# Intensive environmental sampling and whole genome sequence-based characterization of *Listeria* in small and medium sized dairy plants reveal opportunities for simplified and size-appropriate environmental monitoring strategies

**DOI:** 10.1101/2024.01.22.576548

**Authors:** Samantha Bolten, Timothy T. Lott, Robert D. Ralyea, Anika Gianforte, Aljosa Trmcic, Renato H. Orsi, Nicole H. Martin, Martin Wiedmann

## Abstract

Small and medium sized dairy processing plants (SMDPs) may face unique challenges with respect to controlling *Listeria* in their processing environments, e.g., due to limited resources. The aim of this study was to implement and evaluate environmental monitoring programs (EMPs) for *Listeria* control in eight SMDPs in a ∼1-year longitudinal study; this included a comparison of pre-operation (i.e., after cleaning and sanitation and prior to production) and mid-operation (i.e., at least 4 h into production) sampling strategies. Among 2,072 environmental sponge samples collected across all plants, 272 (13%) were positive for *Listeria*. *Listeria* prevalence among pre- and mid-operation samples (15 and 17%, respectively), was not significantly different. Whole genome sequencing (WGS) performed on select isolates to characterize *Listeria* persistence patterns revealed repeated isolation of closely related *Listeria* isolates (i.e., ≤20 high quality single nucleotide polymorphism [hqSNP] differences) in 5/8 plants over >6 months, suggesting *Listeria* persistence and/or re-introduction was relatively common among the SMDPs evaluated here. WGS furthermore showed that for 41 sites where samples collected pre- and mid- operation were positive for *Listeria*, *Listeria* isolates obtained were highly related (i.e., ≤10 hqSNP differences), suggesting that pre-operation sampling alone may be sufficient and more effective for detecting sites of *Listeria* persistence. Importantly, our data also showed that only 1/8 plants showed a significant decrease in *Listeria* prevalence over 1 year, indicating continued challenges with *Listeria* control in at least some SMDPs. We conclude that options for simplified *Listeria* EMP programs (e.g., with a focus on pre-operation sampling, which allows for more rapid identification of likely persistence sites) may be valuable for improved *Listeria* control in SMDPs.

## INTRODUCTION

*Listeria monocytogenes* (LM) is a foodborne pathogen that remains a notable public health concern due to its high case-fatality rate, which has been reported to range from 12.7 to 21% (European Food Safety Authority et al., 2018; Silk et al., 2013). Importantly, LM has been shown to persist for months, and in some cases >12 years, in food processing environments (Y. Chen et al., 2017; Orsi et al., 2008; Vongkamjan et al., 2013), which can increase the risk of cross-contamination to food products (Ferreira et al., 2014). Ready-to-eat (RTE) dairy products (e.g., fluid milk, cheese, ice cream) are a common food vehicle associated with listeriosis outbreaks (Cartwright et al., 2013; Qiu et al., 2021), and, for pasteurized dairy products, past outbreaks have frequently been associated with LM persistence in food processing environments (Acciari et al., 2016; Conrad et al., 2023; Nüesch-Inderbinen et al., 2021). Recently, several listeriosis outbreaks and recalls associated with pasteurized dairy products have been traced back to LM contamination in the processing environment of small and medium sized dairy processing plants (SMDPs) (Centers for Disease Control and Prevention, 2018, 2022; U.S. Food and Drug Administration, 2023), highlighting a need for the development and implementation of strategies that improve *Listeria* control in smaller establishments.

As the presence of non-LM *Listeria* spp. in food processing environments can indicate conditions that might also facilitate the presence of LM (Chapin et al., 2014), regulatory guidance and international standards recommend that food processing plants implement environmental monitoring programs (EMPs), to manage both LM and other *Listeria* spp. (collectively referred to henceforth as “*Listeria*”) in their processing environments (Canadian Food Inspection Agency, 2023; Luber, 2011; U.S. Department of Agriculture Food Safety Inspection Service, 2014; U.S. Food and Drug Administration, 2017). Typical *Listeria* EMPs follow a “seek and destroy” framework, in which a facility will (i) routinely conduct environmental sampling (i.e., collecting sponges or swabs followed by molecular or culture-based *Listeria* detection methods) to “seek” out *Listeria* at discrete sites, and, upon identifying *Listeria* presence in a given site(s), will (ii) perform follow-up activities (e.g., corrective actions, root cause analysis) to “destroy” *Listeria* at that site(s), and also remove or remediate sites of *Listeria* persistence (Malley et al., 2015). However, food processing plants can be very diverse with respect to industry type and size, making it difficult to identify universal standards of appropriate *Listeria* EMP-specific practices (Tompkin, 2002). Some guidance documents recommend that environmental samples be collected mid- operation (e.g., 3 to 4 h into production) for *Listeria* EMPs (Canadian Food Inspection Agency, 2023; U.S. Food and Drug Administration, 2017). This may pose challenges for smaller plants without dedicated food safety personnel, as it could require them to interrupt their production activities and subsequentially increase risk of food safety issues (Magiya, 2023). Given these challenges, SMDPs may benefit from employing more targeted and resource efficient *Listeria* environmental sampling strategies, such as collecting environmental samples pre-operation (i.e., after cleaning and sanitation and prior to a given production cycle), or using passive sampling devices (e.g., Moore swabs in drains) to collect environmental samples mid-operation, as both are less likely to result in production interruption.

Molecular subtyping tools can allow food plants to identify re-occurring subtypes that may be persisting in their food production environments (Beno et al., 2016; Vongkamjan et al., 2013). Whole genome sequencing (WGS) is now typically the preferred molecular subtyping tool for *Listeria* characterization (Jackson et al., 2016), as it can provide valuable insights about *Listeria* transmission and persistence in food processing environments through high-resolution subtyping approaches such as core genome multi-locus sequencing typing (cgMLST) or high quality single nucleotide polymorphism (hqSNP) analysis (Jagadeesan et al., 2019; Sullivan et al., 2022). Additionally, WGS data can be used to screen for high priority genes that may contribute to *Listeria* virulence, as well as tolerance to sanitizers and other environmental stressors (Y. Chen et al., 2020; Daeschel et al., 2022; Kim et al., 2018). Such data can be useful for identifying *Listeria* strains that are potentially of higher priority for food safety interventions.

The purpose of this study was to implement *Listeria* EMPs in eight SMDPs over ∼1 year, and evaluate how *Listeria* EMP implementation impacts the prevalence of *Listeria* in these plants. Additionally, we assessed whether collecting environmental sponge samples pre-operation, and using passive sampling devices (e.g., Moore swabs and tampon swabs) to collect environmental samples mid-operation can represent simplified and effective *Listeria* sampling strategies in SMDPs. Finally, we performed WGS on select *Listeria* isolates obtained in this study to facilitate high-resolution characterization of *Listeria* persistence and transmission patterns in SMDPs.

## MATERIALS AND METHODS

### Study design and *Listeria* EMP implementation

A total of eight SMDPs, located in the Northeastern U.S., and primarily manufacturing either fluid milk (*n*=3), cheese (*n*=2), or ice cream (*n*=3), were conveniently selected for inclusion in this study based on characteristics of the operation and willingness to participate (Table 1). The majority (6/8) of these plants process less than 10 million lb of raw milk annually; we classified these plants as “small dairy processing plants.” The remaining two plants (i.e., plants N and W) process between 10 and 100 million lb of raw milk annually, and were classified as “medium sized dairy processing plants.” Among the small plants enrolled in this study, 5/6 did not have *Listeria* EMPs in place prior to study initiation, while both medium sized plants did have *Listeria* EMPs in place prior to study initiation (Table 1).

**Table 1.** Characteristics of the eight small and medium sized dairy plants (SMDPs) included in the longitudinal study.

| Plant code | Dairy commodity produced | Environment<sup>a</sup> | Estimated annual vol of raw milk processed (lb milk/yr) | EMP<sup>b</sup> in place prior to study initiation | Active ingredients of chemicals used as sanitizers |
| --- | --- | --- | --- | --- | --- |
| CL | Fluid milk | Farmstead | 1,200,000 | No | Peroxyacetic acid |
| CM | Cheese | Standalone | 3,000,000 | No | Quat<sup>c</sup><br>Sodium hypochlorite |
| CN | Ice cream | Standalone | 150,000 | No | Quat<br>Sodium hypochlorite |
| CO | Ice cream | Standalone | 175,000 | No | Peroxyacetic acid<br>Sodium hypochlorite |
| CP | Ice cream | Standalone | 4,000 | No | Quat<br>Sodium hypochlorite |
| CQ | Cheese | Farmstead | 20,000 | Yes | Quat<br>Sodium hypochlorite |
| N | Fluid milk | Standalone | 85,000,000 | Yes | Quat<br>Hydrogen peroxide<br>Sodium hypochlorite |
| W | Fluid milk | Standalone | 15,000,000 | Yes | Sodium hypochlorite |

<sup>a</sup> Farmstead plants are located within 100 m of a dairy farm that provides raw milk for processing.
<sup>b</sup> Refers to whether the plant had a *Listeria* environmental monitoring program (EMP) in place prior to enrollment in this study.
<sup>c</sup> Quaternary ammonium compounds. Includes alkyl-(C8-C18) dimethyl benzyl ammonium chlorides, as well as other compounds represented in the quaternary ammonium compound family.

Upon study initiation, each enrolled plant was visited by at least two members of the research team to facilitate the development of individualized *Listeria* sampling plans, which included development of sampling site lists specific to a given plant. Additionally, during this visit an “initial sampling event” was carried out in which all sites identified on site lists were swabbed using environmental sponges to identify baseline levels of *Listeria* prevalence at the start of the project. For the initial sampling event, environmental sponges were collected at two time points: (i) pre-operation (i.e., after cleaning and sanitation activities were performed, but before the next production cycle) and mid-operation (i.e., at least 4 h into a given production cycle). All pre- and mid-operation environmental sponges were collected within 24 h of each other. For 7/8 enrolled plants, pre- and mid-operation environmental sponges were collected from the same sites, while in one plant (i.e., plant W) distinct sites with similar characteristics (i.e., Zone, site structure) were swabbed pre-operation vs. mid-operation.

Following the initial sampling event, individualized EMP Standard Operating Procedures (SOPs) documents were developed for each enrolled plant. EMP SOPs contained environmental sampling procedures that aligned with FDA’s guidance for controlling *Listeria* in RTE food products (U.S. Food and Drug Administration, 2017). Specifically, plants were instructed to only carry out environmental sampling mid-operation (i.e., at least 4 h into production). Each plant was followed for 11 to 13 months (i.e., ∼1 year) with respect to *Listeria* EMP implementation. During this study period, the research team collected information pertaining to environmental samplings for *Listeria* carried out by plant personnel (referred to as “routine sampling events”) including (i) number of samples collected, (ii) sites that were swabbed, and (iii) test results (i.e., *Listeria* detection). Relevant information on sanitation standard operating procedures (SSOPs) used by plant personnel, including (i) chemicals used for cleaning and sanitation, and (ii) frequency of cleaning and sanitation activities were also collected through informal interviews with plant personnel during the study period. Additionally, the research team provided remote and/or on-site consultations with plant personnel upon request to support *Listeria* control activities (i.e., corrective actions, root cause analysis); during on-site consultations the research team would also collect∼ 6-7 additional environmental sponge samples in “investigational sampling events.”

After ∼1 year of following each plant, a “follow-up sampling event” was carried out by the research team, in which the majority of environmental sponge samples were collected from the same sites that were swabbed in the initial sampling event, to identify changes in *Listeria* prevalence and contamination patterns. In unique instances in which a site sampled in the initial sampling event was no longer available, site substitutions were made in the follow-up sampling event based on the similarity to the initial sampling site.

### Environmental sponge sample collection, processing, and presumptive *Listeria* colony identification for initial, investigational, and follow-up sampling events

The research team collected environmental sponge samples from each enrolled plant in initial, investigational, and follow-up sampling events using sponge kits that contained sponge swabs hydrated with Dey-Engley (DE) broth and sterile gloves (3M, Saint Paul, MN). After collection, all environmental sponges were transported back to Cornell’s Food Safety Laboratory (FSL) on ice, followed by storage at 4°C for up to 48 h before processing.

Environmental sponges were processed using FDA’s bacteriological analytical manual (BAM) method for isolation of *Listeria* from food and environmental samples (Hitchens et al., 2017), with slight modifications. Briefly, 90 mL of buffered *Listeria* enrichment broth (BLEB, BD, Franklin Lakes, NJ) was added to each sponge, followed by stomaching for 60 s at 230 rpm in a Stomacher 400 Circulator (Seward, Basingstoke, UK). After initial incubation for 4 h at 30°C, 360 μL of *Listeria* selective enrichment supplement (LSES, Oxoid, Basingstoke, UK) was added, followed by subsequent incubation at 30°C for up to 48 h. At 24 h and 48 h of incubation, 50 μL of enrichments were streaked for isolation onto (i) modified Oxford agar (MOX, BD) and (ii) *Listeria monocytogenes* chromogenic plating medium (LMCPM, Bisynth International, Itasca, IL), and MOX and LMCPM plates were incubated for 48 h at 30°C and 35°C, respectively.

After incubation, colonies that fit one of the following three categories for appearance on MOX and LMCPM were defined as presumptive *Listeria* colonies: (i) colonies on MOX that showed round and concave morphology and were pewter/dark brown in color were considered presumptive *Listeria* spp. colonies, (ii) colonies on LMCPM that showed round and convex morphology and were turquoise in color were considered presumptive LM or *L. ivanovii* colonies, and (iii) colonies on LMCPM that showed a similar morphology to presumptive LM and *L. ivanovii* colonies, but that lacked the turquoise chroma (resulting from the lack of presence of phosphatidylinositol phospholipase C) were considered presumptive non-LM and non-*L. ivanovii Listeria* spp. colonies. Following presumptive *Listeria* colony identification, up to two presumptive *Listeria* colonies from each category (i.e., i, ii, or iii) were selected for each sample (for a total of up to 6 colonies) and sub-streaked onto non-selective brain heart infusion agar plates (BHIA, BD), followed by incubation at 37°C for 48 h. In the event that presumptive *Listeria* colonies were detected on only one of the selective agars (MOX or LMCPM), up to four presumptive *Listeria* colonies were selected from that selective agar for sub-streaking onto BHIA. Single colonies from sub-streaked plates were frozen down at −80°C in brain heart infusion broth (BHI, BD) containing 15% glycerol for long-term storage.

### Environmental sponge sample collection, processing, and presumptive *Listeria* colony identification for routine sampling events

For all environmental sponge sample collection events carried out by plant personnel during the study period (i.e., routine sampling events), environmental sponges were sent to accredited testing laboratories and processed for *Listeria* detection using proprietary methodologies. Certificates of analyses (COAs) for each routine sampling event were sent to both plant personnel and the research team. If a testing laboratory identified a given environmental sponge as a *Listeria* presumptive positive, the testing lab was instructed to send a portion of the enrichment associated with that presumptive positive to Cornell’s FSL on ice within 7 days after the presumptive positive result was obtained. Upon receipt, enrichments were streaked onto MOX and LMCPM agars and incubated for 48 h at 30°C and 35°C, respectively. After incubation, procedures for presumptive *Listeria* colony identification, sub-streaking of presumptive *Listeria* colonies onto BHIA, and freezing down of presumptive *Listeria* colonies were all carried out as described above.

### Passive sampling of *Listeria* in drains using Moore and tampon swabs

Three SMDPs (i.e., plants CL, CM, and N) were conveniently selected based on willingness to participate in a pilot study that tested the use of passive sampling devices for recovering environmental *Listeria* in drains. For this pilot study, two passive sampling devices were used, including (i) Moore swabs, which represent a swab classically used for environmental surveillance of enteric pathogens in sewage (Moore et al., 1952), and (ii) tampon swabs, which have recently been highlighted as an accessible tool for surveillance of SARS-CoV-2 in wastewater (J. Li et al., 2022). Moore swabs were constructed by folding 3 by 24-inch strips of cheese cloth (VWR, Radnor, PA) 8 times to form an 8-ply square pad, and tying the center with a piece of nylon thread. Tampon swabs were constructed by first removing commercial cotton tampons (Tampax Super unscented) from their applicator, followed by tying a piece of nylon thread to the end of the string attached to the tampon. Individual Moore swabs and tampon swabs were placed in self-sealing sterilization pouches (Thermo Fisher Scientific, Waltham, MA), and autoclaved at 121°C for 20 min. For each plant, 2-5 drains in primary production areas of each plant were identified, and Moore swabs, followed by tampon swabs, were suspended in each drain by nylon thread tied to drain grates ∼3 inches apart from one another. All Moore and tampon swabs were placed in select drains at approximately 30 min to 1 h into a single production cycle. At the end of each plant’s production cycle (∼7-10 h), Moore and tampon swabs were removed from drains, and each swab was aseptically placed into a 55-oz Whirl-Pak bag (Nasco, Fort Atkinson, WI) pre-filled with 10 mL of DE broth. Swabs were transported on ice to Cornell’s FSL, followed by storage at 4°C for up to 24 h before processing. For processing, 90 mL of BLEB was added to Whirl-Pak bags containing either Moore or tampon swabs; all samples were subsequentially processed for *Listeria* detection as described above for environmental sponge samples.

### *Listeria* colony confirmation and *sigB* allelic typing

All presumptive *Listeria* isolates were confirmed as *Listeria* through PCR amplification and Sanger sequencing of a 780-bp fragment of *sigB* as previously described (Sullivan & Wiedmann, 2020). Sanger sequences of the 780-bp *sigB* fragment were queried against the Food Microbe Tracker database (Vangay et al., 2013) to further characterize *Listeria* isolates to the species level, and to assign *sigB* allelic types (ATs) for each isolate.

### Whole genome sequencing

Whole genome sequencing (WGS) was performed on select putatively persistent *Listeria* isolates to characterize the longitudinal persistence of *Listeria* in SMDPs. Putative persistence was identified using *sigB* AT data, such that if a given *sigB* AT of *Listeria* was isolated from a given plant in at least 2 samplings that were conducted at least 7 days apart, then that *sigB* AT was considered putatively persistent in the plant. One *Listeria* isolate from each sample that contained a putatively persistent *sigB* AT for a given plant was randomly selected for sequencing. Additionally, if a given *sigB* AT was only identified in samplings conducted <7 days apart, but was identified at the same site for both pre-operation and mid-operation sampling times (i.e., in initial and follow-up sampling events), then that *sigB* AT was also considered putatively persistent, and one isolate per sample was randomly selected for sequencing.

Single colonies of selected isolates were inoculated into 5 mL BHI, incubated for 15-18 h at 37°C, and DNA from cultures was extracted using the QiAmp DNA minikit (Qiagen, Germantown, MD) according to manufacturer instructions. Sequencing of genomic DNA was performed using the Illumina NextSeq 500 platform (Illumina, Inc., San Diego, CA) with a maximum read length of 2 x 150 bp. Raw reads were trimmed using Trimmomatic v. 0.39, and contigs were evaluated and filtered based on quality using FastQC v. 0.11.8 (Bolger et al., 2014). Filtered contigs were assembled using SPAdes v. 3.15.4 using careful mode (Bankevich et al., 2012). Quality control of genome assemblies was performed using QUAST v. 5.0.2 (Gurevich et al., 2013), and average coverage was determined using SAMtools v. 1.14 (H. Li et al., 2009). Assemblies with a genome-wide average coverage greater than 50x were included in genomic analysis. Contigs smaller than 500 bp, and with an average coverage <6x were removed. All genome assemblies are publicly available on the Cornell University eCommons repository: https://doi.org/10.7298/ah3b-ew05 (Bolten et al., 2023), and quality metrics for all genome assemblies are provided in Supplementary Table 1.

### High-quality SNP analysis

All isolates were sorted into groups based on *sigB* AT, and a reference-free SNP analysis using kSNP3 v. 3.1 was performed to determine clusters for hqSNP analysis (Gardner et al., 2015). Isolates <100 kSNPs apart identified based on kSNP3 analysis were then analyzed using the Center for Food Safety and Applied Nutrition (CFSAN) SNP pipeline v. 2.2.1 to identify any preserved hqSNPs existing between groups of isolates (Davis et al., 2015). The reference assembly for the CFSAN SNP pipeline was selected as previously described (Sullivan et al., 2022). After hqSNP analysis was performed for groups of isolates with <100 kSNP differences, isolates that showed >50 pairwise hqSNP differences from all other isolates in the group were removed, and the group was re-examined to obtain “hqSNP clusters” which only included isolates that showed ≤50 pairwise hqSNP differences from at least one other isolate in the cluster. Maximum likelihood phylogenetic trees based on hqSNP data were created using RAxML (v. 8.2.12) with 1,000 bootstraps and the GTRCAT nucleotide substitution model (Stamatakis, 2014).

### Gene presence or absence queries

The nucleotide sequences for select *Listeria* pathogenicity islands (LIPI-3 and LIPI-4), stress survival islets (SSI-1 and SSI-2), and sanitizer and metal tolerance genes (*bcrABC*, *emrE*, *qacA*, *qacH* [*emrC*], *cadAC*) were downloaded from the Pasteur Institute (https://bigsdb.pasteur.fr/cgi-bin/bigsdb/bigsdb.pl?db=pubmlst_listeria_seqdef&page=downloadAlleles on May 26^th^, 2023. All sequences were used to create a local nucleotide BLAST database, and the database was searched against *Listeria* genome assemblies using blastn and default parameters. Matches with >90% sequence identity and >90% query coverage indicated the presence of a gene in the respective isolate’s genome. Additionally, identification of SNPs leading to premature stop codons (PMSCs) in *inlA* was performed as previously described (Harrand et al., 2020). For LM genome assemblies, sequence types (ST), and clonal complexes (CC) were assigned based on *in silico* MLST subtyping of seven MLST loci (i.e., *abcZ*, *bglA*, *cat*, *dapE*, *dat*, *ldh*, *lhkA*) (Ragon et al., 2008), available at the BIGsdb-Lm (Moura et al., 2016).

### Ontology-based classification of site structures

The structures of sites that were swabbed in this study were captured from unique free-text sampling site descriptions based on a previously reported schema (Feng et al., 2023). Briefly, terms used to indicate the structure that was swabbed in each free-text sampling site description were identified and queried against the Ontobee database (https://ontobee.org) to match those terms with existing ontologies procured from databases including the Environmental Ontology (ENVO), the Phenotype and Trait Ontology (PATO), and National Cancer Institute Thesaurus OBO Edition (NCIT). In most cases, the structure of a site could be captured based on one ontology (e.g., drain, ENVO:01000924), while in some specific cases, two or more ontologies were used to capture the appropriate level of granularity associated with the site’s structure (e.g., pallet jack [NCIT:C50012], wheels [ENVO:03501261]), as described in (Feng et al., 2023).

### Statistical analysis

All data analyses were performed in R version 4.0.2. Generalized linear regression models were used to assess whether there was a statistically significant difference in the probability of a given sample testing positive for *Listeria* between initial and follow-up sampling events within a given dairy plant. Additionally, a generalized linear mixed effects regression model using data from the initial and follow-up sampling events was used to assess the effect of sampling time (i.e., pre-operation vs. mid-operation) and Zone (i.e., Zone 2, 3, or 4) on the probability of a given sample testing positive for *Listeria* across all dairy plants. For this analysis, sampling event, sampling time and Zone were considered fixed effects, and dairy plant nested with site was considered a random effect. *Post-hoc* analyses of both (i) generalized linear regression models and (ii) generalized linear mixed effects regression models were carried out using the emmeans package in R (Lenth, 2019).

## RESULTS

### *Listeria* prevalence varied across different SMDPs over the 1-year study period

Environmental sponge samples from eight enrolled SMDPs were collected by members of the research team in two separate sampling events: (i) an initial sampling event prior to *Listeria* EMP implementation, and (ii) a follow-up sampling event ∼1 year after *Listeria* EMP implementation (Table 2). For the initial sampling event, a total of 561 environmental sponges were collected (range of 42 to 116 sponges per plant), and for the follow-up sampling event, a total of 558 environmental sponges were collected (range of 44 to 129 sponges per plant) (Table 3). Additionally, plant personnel collected environmental sponge samples in (iii) routine sampling events throughout the ∼1 year study period. For routine sampling events, a total of 927 environmental sponges were collected across all plants, with a median of 28 (range of 0 to 562) environmental sponges collected per plant. Finally, for four plants (i.e., plants CL, CM, CN, and CO) the research team also collected environmental sponge samples in (iv) investigational sampling events during on-site consultations that were requested by the plant (Table 2). For investigational sampling events, a total of 26 environmental sponge samples were collected (range of 6 to 7 sponges per plant).

**Table 2.** Timeline of sampling events in the longitudinal study.

| Plant Code | Sampling time<sup>c</sup> | Initial sampling event date | Follow-up sampling event date | Routine sampling events<sup>a</sup> |  | Investigational sampling events<sup>b</sup> |  |
| --- | --- | --- | --- | --- | --- | --- | --- |
|  |  |  |  | No. of events | Year: Julian week in which samples were collected<sup>d</sup> | No. of events | Year: Julian week in which samples were collected |
| CL | Pre-operation | 2/14/2021 | 2/21/2022 | 2 | 2021: 33, 44 | 1 | 2021: 45 |
|  | Mid-operation | 2/15/2021 | 2/21/2022 |  |  |  |  |
| CM | Pre-operation | 2/15/2021 | 2/16/2022 | 4 | 2021: 18, 23, 32 | 1 | 2021: 44 |
|  | Mid-operation | 2/16/2021 | 2/15/2022 |  | 2022: 1 |  |  |
| CN | Pre-operation | 2/22/2021 | 3/7/2022 | 2 | 2021: 18, 34 | 1 | 2021: 43 |
|  | Mid-operation | 2/22/2021 | 3/7/2022 |  |  |  |  |
| CO | Pre-operation | 2/21/2021 | 1/25/2022 | 2 | 2021: 25, 44 | 1 | 2021: 45 |
|  | Mid-operation | 2/22/2021 | 1/25/2022 |  |  |  |  |
| CP | Pre-operation | 2/10/2021 | 3/31/2022 | 0 | NA<sup>e</sup> | 0 | NA |
|  | Mid-operation | 2/10/2021 | 3/30/2022 |  |  |  |  |
| CQ | Pre-operation | 6/17/2021 | 6/3/2022 | 9 | 2021: 31, 35, 38, 44, 46 | 0 | NA |
|  | Mid-operation | 6/17/2021 | 6/2/2022 |  | 2022: 1, 5, 14, 18 |  |  |
| N | Pre-operation | 3/18/2021 | 3/22/2022 | 50 | 2021: 16-22, 24-51, 52<sup>f</sup> | 0 | NA |
|  | Mid-operation | 3/17/2021 | 3/21/2022 |  | 2022: 1-2, 3<sup>f</sup>, 4-6, 7<sup>f</sup>, 8-11 |  |  |
| W | Pre-operation | 9/23/2020 | 10/12/2021 | 22 | 2020: 40, 42, 44-47, 50 | 0 | NA |
|  | Mid-operation | 9/23/2020 | 10/12/2021 |  | 2021: 1, 3, 4<sup>f</sup>, 5, 8, 9, 11, 13-15, 17, 19, 27, 31, 37, 39 |  |  |

<sup>a</sup> Routine sampling events were carried out by personnel from each dairy plant throughout the ~1 year study period. All dairy plant personnel were instructed to perform routine sampling mid-operation.
<sup>b</sup> Investigational sampling events were carried out by the research team during optional on-site consultations that were requested by a given dairy plant. All investigational samples were collected mid- operation.
<sup>c</sup> In initial and follow-up sampling events, environmental sponge samples were collected both pre- operation (i.e., after cleaning and sanitation but before the next production cycle) and mid-operation (i.e., at least 4 h into a given production cycle).
<sup>d</sup> Each Julian week corresponds to 7-Julian dates in an ordinal 365-day year (i.e., Jan 1 to 7 represents Julian week 1)
(http://www.krisweb.com/biblio/trinity_cdfg_cdfg_xxxx_monitoring/9293mon/fta/chp2/app1.pdf)
<sup>e</sup> Not available as routine or investigational sampling events were not carried out.
<sup>f</sup> Indicates Julian weeks in which two routine sampling events were carried out by a given dairy plant.

**Table 3.** Number of environmental sponge samples that were positive for *Listeria* across different sampling events.

| Plant code | Initial sampling event<sup>a</sup> |  | Routine sampling events<sup>b</sup> |  | Investigational sampling event<sup>c</sup> |  | Follow-up sampling event<sup>a</sup> |  | All samplings<sup>d</sup> |  |
| --- | --- | --- | --- | --- | --- | --- | --- | --- | --- | --- |
|  | No. of positive samples (%) | No. of samples collected | No. of positive samples (%) | No. of samples collected | No. of positive samples (%) | No. of samples collected | No. of positive samples (%) | No. of samples collected | No. of positive samples (%) | No. of samples collected |
| CL | 15 (30) | 50 | 4 (40) | 10 | 4 (57) | 7 | 13 (25) | 52 | 36 (30) | 119 |
| CM | 2 (2.7) | 73 | 0 (0) | 37 | 0 (0) | 7 | 2 (2.7) | 74 | 4 (2.1) | 191 |
| CN | 33 (63) | 52 | 8 (80) | 10 | 2 (33) | 6 | 8 (15) | 54 | 51 (42) | 122 |
| CO | 4 (9.5) | 42 | 6 (32) | 19 | 3 (50) | 6 | 9 (20) | 44 | 22 (20) | 111 |
| CP | 2 (2.1) | 94 | — | 0<sup>e</sup> | — | 0 | 25 (36) | 70 | 27 (16) | 164 |
| CQ | 4 (7.5) | 53 | 4 (11) | 36 | — | 0 | 8 (15) | 54 | 16 (11) | 143 |
| N | 1 (0.9) | 116 | 17 (3.0) | 562 | — | 0 | 8 (6.2) | 129 | 26 (3.2) | 807 |
| W | 25 (31) | 81 | 47 (19) | 253 | — | 0 | 18 (22) | 81 | 90 (22) | 415 |
| All plants | 86 (15) | 561 | 86 (9.3) | 927 | 9 (35) | 26 | 91 (16) | 558 | 272 (13) | 2072 |

<sup>a</sup> Refers to data collected in a single sampling event (i.e., initial or follow-up sampling event) that was carried out by the research team over two sampling times (i.e., pre-operation and mid-operation).
<sup>b</sup> Refers to data collected in one or more routine sampling events that were carried out by plant personnel from each dairy plant throughout the ~1 year study period.
<sup>c</sup> Refers to data collected in investigational sampling events carried out by the research team during on-site consultations that were requested by the plant throughout the ~1 year study period.
<sup>d</sup> Represents the total number of environmental sponge samples collected in a given dairy plant across all sampling events (i.e., initial, routine, investigational, and follow-up sampling events).
<sup>e</sup> Plant CP did not collect any routine environmental samples across the 1-year study period.

Overall, across all sampling events and plants, a total of 2,072 environmental sponges were collected (111 to 807 sponges per plant; Table 3). Of these 2,072 environmental sponges, 272 (13%) tested positive for *Listeria*. *Listeria* prevalence varied widely within each plant, with plants CN (51/122, 42%) and CL (36/119, 30%) showing the highest overall *Listeria* prevalence across all samplings, and plants CM (4/191, 2.1%) and N (26/807, 3.2%) showing the lowest overall *Listeria* prevalence across all samplings.

### The impact of *Listeria* EMP implementation on *Listeria* prevalence differed among plants

Across all plants, the number of *Listeria* positive environmental sponges detected in the initial sampling event (86/561, 15%) was numerically similar to the number of positive environmental sponges detected in the follow-up sampling event (91/558, 16%) (Table 3). Generalized linear regression analysis showed that, for 6/8 plants, the probability of detecting *Listeria* from environmental sponges collected in the follow-up sampling event was not significantly different from the probability of detecting *Listeria* in the initial sampling event (Figure 1). Exceptions included plant CN, in which environmental sponges were significantly less likely to test positive for *Listeria* in the follow-up sampling event compared to the initial sampling event (p<0.05), and plant CP, in which environmental sponges were significantly more likely to test positive for *Listeria* in the follow-up sampling event compared to the initial sampling event (p<0.05). Importantly, plant CP was the only plant enrolled in this study that did not carry out any routine sampling events throughout the 1-year study period (Table 2), which anecdotally suggests that a lack of routine sampling for *Listeria* may impede overall *Listeria* control in SMDPs.

**Figure 1.**
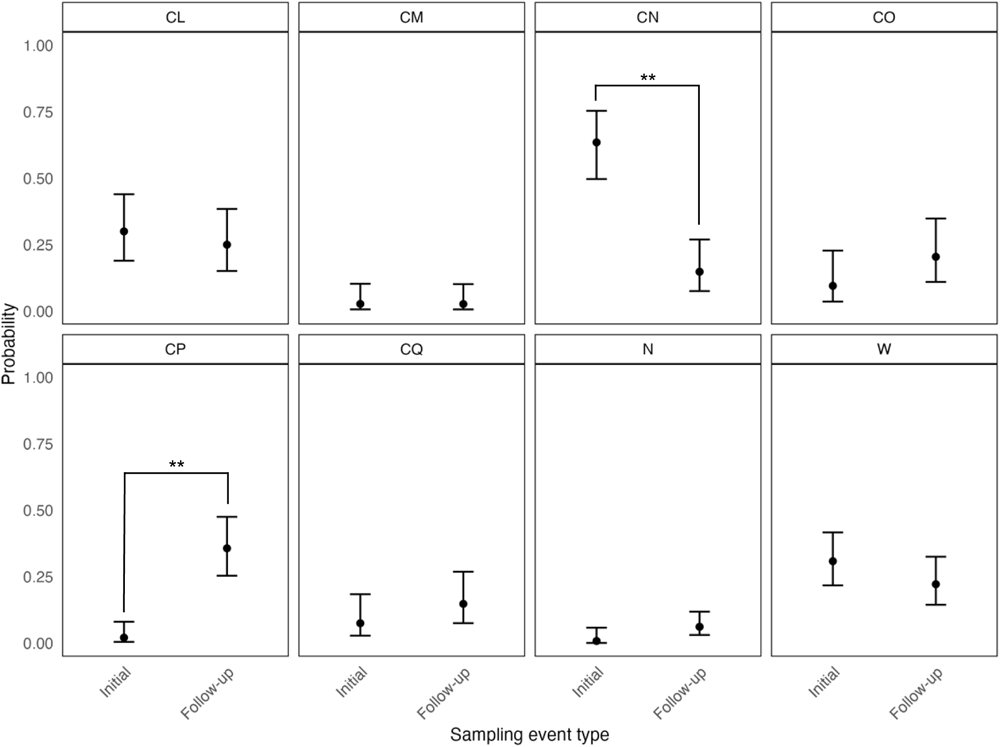
Probability of an environmental sponge sample testing positive for *Listeria* in small and medium sized dairy plants (SMDPs) across initial and follow-up sampling events. Data includes samples collected at both pre-operation and mid-operation sampling times. Error bars represent 95% confidence intervals derived from generalized linear regression models generated for each plant. Within the same panel, pairwise comparisons of model estimates between sampling events was performed using Tukey’s honestly significant difference (HSD) *post hoc* test. Significance codes: “**” represents *P*<0.001.

### Pre-operation sampling yielded similar percentage of *Listeria* positive samples compared to mid-operation sampling

In both the initial and follow-up sampling events, the same or similar (in the case of plant W) sites were swabbed with environmental sponges at two different sampling times: pre-operation (i.e., after cleaning and sanitation, but before the next production cycle), and mid-operation (i.e., at least 4 h into production), to assess for the impact of sampling time on *Listeria* detection in SMDPs. Overall, across both initial and follow-up sampling events for all plants, detection of *Listeria* positive sponges collected pre-operation (84/564, 15%), was numerically similar to those collected mid-operation (93/555, 17%) (Figure 2). Furthermore, generalized linear mixed effects regression analysis revealed that the odds of a given environmental sponge testing positive for *Listeria* was not significantly associated with sampling event (p=0.54) or sampling time (p=0.19). However, the odds of an environmental sponge testing positive for *Listeria* was significantly associated with the Zone (i.e., Zone 2, 3, or 4) that was swabbed (p<0.05). In particular, the odds of a given environmental sponge testing positive for *Listeria* were significantly higher if the site that was swabbed represented either a Zone 3 (0.15) or Zone 4 (0.18) site, compared to Zone 2 (0.012) (p<0.05). Furthermore, across both initial and follow-up sampling events, *Listeria* positive samples from Zone 2 sites were only obtained from 3 out of 8 SMDPs (i.e., plants CL, CP, and W) (Figure 2), suggesting that the majority of SMDPs evaluated here were better able to control *Listeria* in Zone 2 sites compared to Zones 3 and 4 sites.

**Figure 2.**
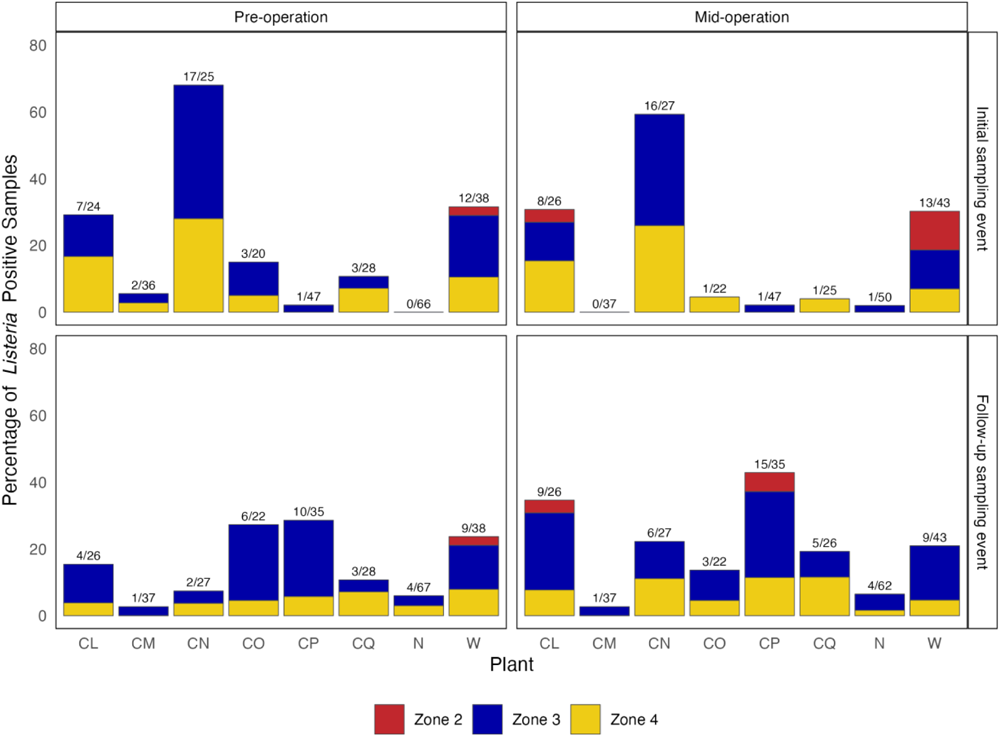
Percentage of *Listeria* positive environmental sponge samples collected either pre-or mid-operation in initial and follow-up sampling events. Different colors within each bar represent the percentage of *Listeria* positive samples by Zone (i.e., Zone 2, 3, or 4). Fractions at the top of each bar indicate the total number of positive samples obtained/total number of samples that were collected for a given sampling event and sampling time.

### Moore and tampon swabs placed in drains yielded similar detection of *Listeria* when compared to environmental sponges

During the follow-up sampling events for plants CL, CM, and N, Moore and tampon swabs were placed in 2 to 5 conveniently selected drains for the duration of a single production cycle to evaluate the ability of these passive sampling devices to detect *Listeria* in the drains of SMDPs. In total, 12 drain sites were evaluated for *Listeria* detection using Moore and tampon swabs (*n*=2, *n*=5, and *n*=5 for plants CL, CM, and N, respectively), of which 9 also represented sites that were also sampled using environmental sponges in the follow-up sampling event (*n*=2, *n*=3, and *n*=3 for plants CL, CM, and N, respectively). Overall, 2/12 (17%) Moore swabs, and 2/12 (17%) tampon swabs yielded *Listeria* positive results; all positive Moore/tampon swabs were obtained from plant CL at drain sites CL1 and CL2 (Table 4). Environmental sponges collected mid-operation from drain sites CL1 and CL2 during the same production cycle also yielded *Listeria* positive results, supporting that Moore/tampon swabs and environmental sponges showed comparable ability to detect *Listeria* at these drain sites. Interestingly, *Listeria* was not detected from environmental sponges collected pre-operation from drain sites CL1 and CL2 in the same follow-up sampling event, which suggests *Listeria* introduction at these drain sites during the production cycle, rather than *Listeria* persistence at these drain sites.

**Table 4.** Detection of *Listeria* in Moore and tampon swabs placed in drains throughout a single production cycle in the follow-up sampling event for plants CL, CM, and N.

| Drain No.<br>(site ID) | *Listeria* detection result using |  |  |  |
| --- | --- | --- | --- | --- |
|  | Moore swab<sup>a</sup> | tampon swab<sup>a</sup> | sponge<sup>b</sup>, mid-operation | sponge<sup>b</sup>, pre-operation |
| **Plant CL** |  |  |  |  |
| Drain 1 (CL1) | Positive | Positive | Positive | Negative |
| Drain 2 (CL2) | Positive | Positive | Positive | Negative |
| **Plant CM** |  |  |  |  |
| Drain 1 (CM3) | Negative | Negative | Negative | Negative |
| Drain 2 (CM1) | Negative | Negative | Negative | Negative |
| Drain 3 (CM21) | Negative | Negative | Negative | Negative |
| Drain 4 (NA<sup>c</sup>) | Negative | Negative | —<sup>c</sup> | — |
| Drain 5 (NA) | Negative | Negative | — | — |
| **Plant N** |  |  |  |  |
| Drain 1 (N12) | Negative | Negative | Negative | Negative |
| Drain 2 (N22) | Negative | Negative | Negative | Negative |
| Drain 3 (N38) | Negative | Negative | Negative | Negative |
| Drain 4 (NA) | Negative | Negative | — | — |
| Drain 5 (NA) | Negative | Negative | — | — |

<sup>a</sup> All Moore and tampon swabs were collected during the follow-up sampling event on 2/21/2022, 2/16/2022, and 3/22/2022, for plants CL, CM, and N, respectively.
<sup>b</sup> All sponge samples were collected in the follow-up sampling event. For dates of sampling see Table 2.
<sup>c</sup> Not applicable as the drain in which Moore and tampon swabs were collected did not represent a site in which environmental sponge samples were collected in the follow-up sampling event.

### Repeated isolation of *sigB* ATs across multiple samplings was observed for 7/8 SMDPs

Across all sampling events, positive environmental samples from sponges (*n*=272), Moore swabs (*n*=2), and tampon swabs (*n*=2) yielded a total of six different species of *Listeria*, including LM, *L. grayi*, *L. innocua*, *L. newyorkensis*, *L. seeligeri*, and *L. welshimeri*. Select species were isolated from multiple plants, including *L. innocua* (isolated from all 8 plants), as well as LM (6 plants), *L. seeligeri* (4 plants), and *L. welshimeri* (3 plants) (Table 5). *L. grayi* and *L. newyorkensis* were only isolated from a single plant (plant W and plant CO, respectively).

**Table 5.** Number of environmental sponge, Moore swab, and tampon swab samples that tested positive for a given species of *Listeria*.

| Plant code | No. of samples collected<sup>a</sup> | No. of positive samples | No. of samples positive for a given species of *Listeria* (%)<sup>b</sup> |  |  |  |  |  |
| --- | --- | --- | --- | --- | --- | --- | --- | --- |
|  |  |  | LM<sup>c</sup> | *L. innocua* | *L. seeligeri* | *L. welshimeri* | *L. grayi* | *L. newyorkensis* |
| CL | 123 | 40<sup>d</sup> | 6 (4.9) | 36 (29) | 2 (1.6) | 0 (0) | 0 (0) | 0 (0) |
| CM | 201 | 4 | 0 (0) | 4 (1.9) | 0 (0) | 0 (0) | 0 (0) | 0 (0) |
| CN | 122 | 51 | 0 (0) | 51 (42) | 0 (0) | 0 (0) | 0 (0) | 0 (0) |
| CO | 111 | 22 | 1 (0.9) | 3 (2.7) | 3 (2.7) | 1 (0.9) | 0 (0) | 17 (15) |
| CP | 164 | 27 | 2 (1.2) | 25 (15) | 0 (0) | 0 (0) | 0 (0) | 0 (0) |
| CQ | 143 | 16 | 7 (4.9) | 4 (2.8) | 9 (6.3) | 0 (0) | 0 (0) | 0 (0) |
| N | 817 | 26 | 5 (0.6) | 21 (2.6) | 0 (0) | 3 (0.4) | 0 (0) | 0 (0) |
| W | 415 | 90 | 75 (18) | 26 (6.2) | 20 (4.8) | 1 (0.2) | 4 (1.0) | 0 (0) |

<sup>a</sup> Represents the total number of environmental sponge, Moore swab, and tampon swab samples collected for a given dairy plant across all sampling events (i.e., initial, routine, investigational, and follow-up sampling events), and sampling times (i.e., pre-operation and mid-operation).
<sup>b</sup> Percentage was calculated with respect to the total number of samples collected from each plant.
<sup>c</sup> *L. monocytogenes*.
<sup>d</sup> In some cases, multiple species were identified from the same sample. Two different species were isolated from 30, 4, 3, 3, and 2 samples from plants W, CL, CO, N, and CQ, respectively. Three different species were isolated from 3 samples from plant W, and 1 sample from plant CQ.

Of the 276 environmental samples (i.e., sponges, Moore swabs, tampon swabs) that tested positive for *Listeria*, 1,071 isolates were obtained and further characterized into distinct *sigB* ATs. Of the 1,071 isolates, 364 were classified as representative isolates, which was defined as an isolate from a given sample with a distinct *sigB* AT. In total, 36 distinct *sigB* ATs were identified across plants, with a range of 1-4 *sigB* ATs being isolated from a given sample (median: 1 *sigB* AT per sample); a total of 67/276 (24%) environmental samples yielded more than 1 *sigB* AT (Figure 3).

**Figure 3.**
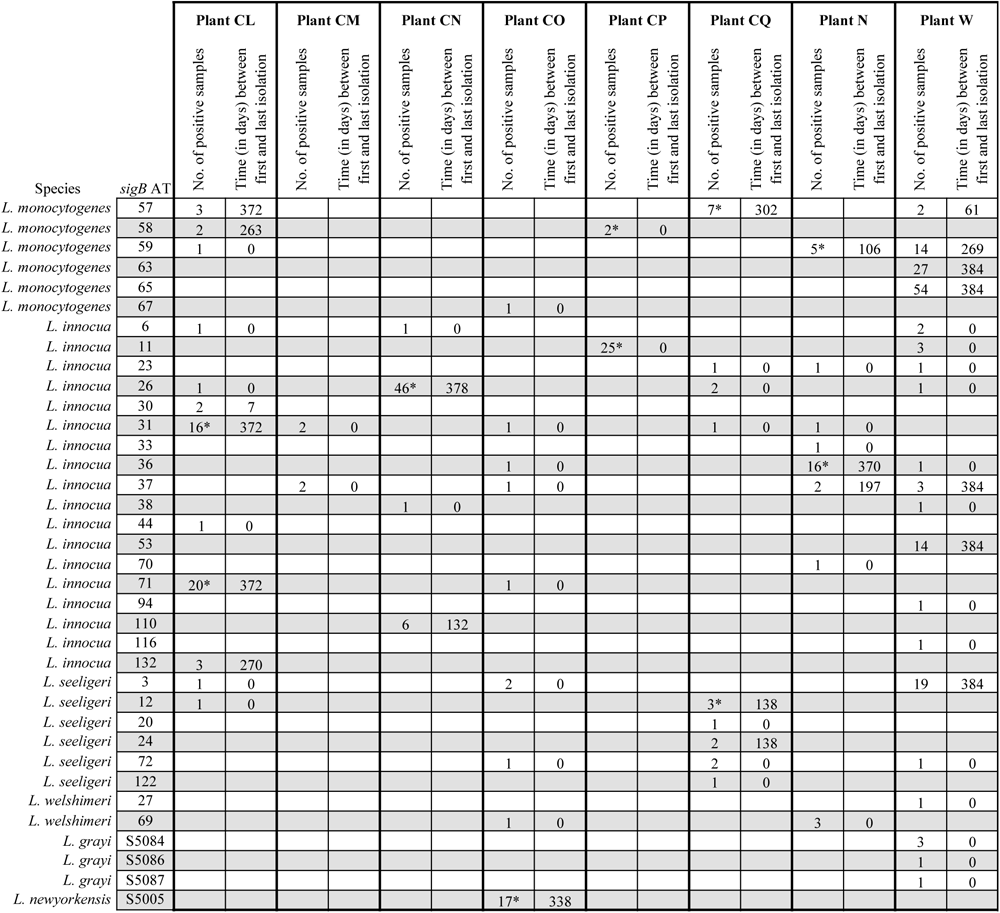
Occurrence of different *Listeria sigB* allelic types (ATs) in environmental sponge, Moore swab, and tampon swab samples collected from small and medium sized dairy processing plants (SMDPs) over the 1-year study period. A “*” indicates that a given *sigB* AT was obtained from the same site at both pre-operation and mid-operation sampling times during a given sampling event (i.e., initial and/or follow-up sampling).

For 283/364 representative isolates from 6/8 dairy plants (i.e., plants CL, CN, CO, CQ, N, and W), the same *sigB* AT was repeatedly isolated from the same plant over ≥7 days, suggesting putative persistence (Figure 3). In addition, 20/364 of the representative isolates, all obtained from plant CP, represented repeated isolation of the same *sigB* AT at the same site when sampled at both pre- and mid- operation sampling times on the same date (i.e., 2/2 isolates representing LM AT 58, and 18/25 isolates representing *L. innocua* AT 11), also suggesting putative persistence. Thus, a total of 303 representative isolates, across all plants except plant CM, represented a putatively persistent *Listeria sigB* AT. One isolate representing each of these putatively persistent *sigB* ATs was randomly selected from each sample and subjected to WGS characterization. A list of the metadata associated with each WGS-characterized isolate, including species, *sigB* AT, sampling time, month and year of isolation, and hqSNP cluster assignment, is provided in Supplementary Table 2.

### Over 90% of WGS-characterized *Listeria* isolates could be grouped into hqSNP clusters

The 303 putatively persistent *Listeria* isolates characterized through WGS represented *L. innocua* (*n*=146), LM (*n*=116), *L. seeligeri* (*n*=24), and *L. newyorkensis* (*n*=17) (Supplementary Table 2). As expected, given our selection criteria, the majority of *Listeria* isolates sequenced here could be grouped into hqSNP clusters (286/303, 94%), while the remaining isolates represented singletons (17/303, 6%) that showed >50 hqSNP differences from all other WGS-characterized *Listeria* isolates. A total of 25 hqSNP clusters were identified, with the number of isolates represented in a given cluster ranging from 2 to 54 (Table 6). All hqSNP clusters were comprised of isolates representing a single plant, with plant W showing the highest number of hqSNP clusters (*n*=8), followed by plant CL (*n*=5), and plant N (*n*=4). As previous studies have suggested using various SNP cut-offs, such as 20 SNPs (Pightling et al., 2018; Wang et al., 2018), or 10 SNPs (European Centre for Disease Prevention and Control, 2016), to indicate whether two or more isolates originated from the same source population, *Listeria* isolates examined in this study were further classified with respect to 5 arbitrary hqSNP cut-offs: (i) distantly related (>50 hqSNP differences), (ii) loosely related (>21 but ≤50 hqSNP differences), (iii) closely related (>10 but ≤20 hqSNP differences), (iv) highly related (>2 but ≤10 hqSNP differences), or (v) nearly identical (0-2 hqSNP differences). With respect to these cut-offs, all isolates represented in 1, 5, 5, 12, and 2 hqSNP clusters were classified as distantly related, loosely related, closely related, highly related, and nearly identical, respectively (Table 6).

**Table 6.** Characteristics of *Listeria* isolates represented in hqSNP clusters.

| Cluster no. | Plant code | *sigB* AT<sup>a</sup> | No. of isolates | Time (in months) between first and last isolation | hqSNP range | Relatedness classification<sup>b</sup> |
| --- | --- | --- | --- | --- | --- | --- |
| ***L. monocytogenes* clusters** |  |  |  |  |  |  |
| 1 | CL | 58 | 2 | 8.6 | 14 | Closely related |
| 2 | CP | 58 | 2 | 0<sup>c</sup> | 0 | Nearly identical |
| 3 | CQ | 57 | 6 | 0 | 0-2 | Nearly identical |
| 4 | N | 59 | 3 | 2.8 | 7-12 | Closely related |
| 5 | N | 59 | 2 | 0 | 7 | Highly related |
| 6 | W | 59 | 14 | 8.8 | 0-10 | Highly related |
| 7 | W | 63 | 27 | 12.6 | 1-25 | Loosely related |
| 8 | W | 65 | 54 | 12.6 | 0-42 | Loosely related |
| ***L. innocua* clusters** |  |  |  |  |  |  |
| 9 | CL | 30 | 2 | 0.2 | 4 | Highly related |
| 10 | CL | 31 | 14 | 12.2 | 0-53 | Distantly related |
| 11 | CL | 71 | 20 | 12.2 | 0-5 | Highly related |
| 12 | CL | 132 | 3 | 8.9 | 5-9 | Highly related |
| 13 | CN | 26 | 46 | 12.4 | 0-13 | Closely related |
| 14 | CN | 110 | 5 | 4.3 | 1-7 | Highly related |
| 15 | CP | 11 | 17 | 0 | 0-4 | Highly related |
| 16 | N | 36 | 14 | 12.1 | 0-18 | Closely related |
| 17 | N | 36 | 2 | 0 | 10 | Highly related |
| 18 | W | 53 | 11 | 12.6 | 1-21 | Loosely related |
| 19 | W | 53 | 3 | 0 | 4-34 | Loosely related |
| ***L. seeligeri* clusters** |  |  |  |  |  |  |
| 20 | CQ | 12 | 2 | 0 | 9 | Highly related |
| 21 | CQ | 24 | 2 | 4.5 | 18 | Closely related |
| 22 | W | 3 | 5 | 10.4 | 3-7 | Highly related |
| 23 | W | 3 | 9 | 12.6 | 0-35 | Loosely related |
| 24 | W | 3 | 4 | 12.6 | 3-6 | Highly related |
| ***L. newyorkensis* clusters** |  |  |  |  |  |  |
| 25 | CO | S5005 | 17 | 11.1 | 0-5 | Highly related |

<sup>a</sup> *sigB* allelic type.
<sup>b</sup> Clusters were classified as either (i) distantly related (>50 hqSNP differences), (ii) loosely related (>21 but ≤50 hqSNP differences), (iii) closely related (>10 but ≤20 hqSNP differences), (iv) highly related (>2 but ≤10 hqSNP differences), or (v) nearly identical (0-2 hqSNP differences).
<sup>c</sup> For isolates obtained from samples collected within 24 h of one another (i.e., isolates collected at pre- operation and mid-operation sampling times in either the initial or follow-up sampling event), months between first and last isolation was reported as “0”.

Singletons were identified across all plants (range of 1-6 singletons per plant), except for plant CO, in which all (17/17) *L. newyorkensis* isolates sequenced were represented in cluster 25. Overall, the majority (12/17, 71%) of singletons showed >100 kSNPs from all other isolates characterized through WGS, suggesting that they were likely transient in their respective plants (Supplementary Table 3). Exceptions included FSL B10-0031 and FSL R12-1459 from plant W, which showed 57 pairwise hqSNP differences from one another and were isolated >2 months apart, and FSL B10-0396 and FSL L8-0895 from plant CL, which showed 80 pairwise hqSNP differences from one another and were isolated >12 months apart. Additionally, singleton FSL B10-0243 from plant CL showed 73-78 hqSNP differences from all isolates represented in plant CL cluster 10, and was isolated ∼8 months after the first isolation of cluster 10.

### Persistence/re-introduction of *Listeria* isolates over >6 months was observed in five plants

Overall, more than half (14/25, 56%) of hqSNP clusters contained isolates that were obtained >6 months apart, with the majority of these hqSNP clusters being obtained from plant W (*n*=7), followed by plant CL (*n*=4), and plants CN, CO, and N (all *n*=1) (Table 6). Of these 14 hqSNP clusters, the majority (9/14, 64%) either contained isolates that were closely related (*n*=3) or highly related (*n*=6), which suggests that they likely share a common source that could be either in the plant, which would indicate persistence, or outside of the plant, which would indicate re-introduction into the plant. Importantly, although the remaining 5/14 hqSNP clusters that showed re-isolation over >6 months contained at least one isolate that was loosely or distantly related (>21 hqSNP differences) compared to all other isolates in the hqSNP cluster (i.e., clusters 7, 8, 10, 18, and 23), in many cases these hqSNP clusters contained closely or highly related monophyletic subclusters that were isolated >6 months apart. For example, plant W LM cluster 7 contained 3 closely or highly related subclusters that were isolated >6 months apart (Figure 4A), and plant W *L. seeligeri* cluster 23 contained 2 highly related monophyletic subclusters that were isolated >6 months apart (Figure 5A). This may suggest that these hqSNP clusters have been persisting in these plants for several years, allowing them time to evolve and accumulate a high number of SNPs. Interestingly, these closely/highly related cluster 7 and cluster 23 subclusters were isolated in at least one sampling from site W65, a drain in the filling and casing area of plant W (Figure 4B, 5B), which may represent a potential long-term harborage site for these clusters.

**Figure 4.**
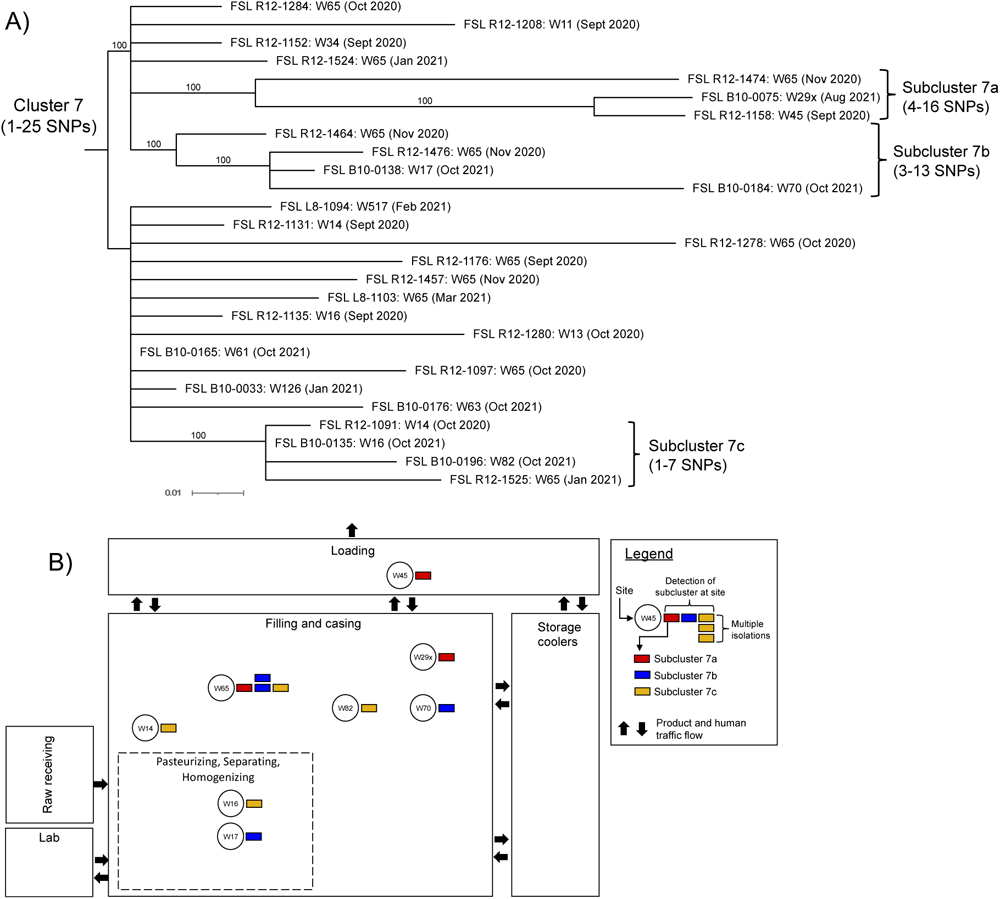
Maximum likelihood phylogenetic tree of plant W *L. monocytogenes* cluster 7 and its subclusters (A) and map of the location in which subclusters were identified in the plant (B). For (A), each node represents isolate ID (i.e., FSL R12-1284), site number (i.e., W65), and date (month and year) in which a given isolate was obtained. hqSNP data from cluster 7 was used to create the maximum likelihood phylogenetic tree using RAxML (v 8.2.12) with 1,000 bootstraps and the GTRCAT nucleotide substitution model. Only bootstrap values >70% are presented in the tree, and the tree is rooted at its midpoint. The scale at the bottom of the phylogenetic tree indicates genetic distance. For (B), a colored bar indicates isolation of a given subcluster, and multiple bars of the same color are shown for a given site to indicate the number of samplings when a subcluster was isolated.

**Figure 5.**
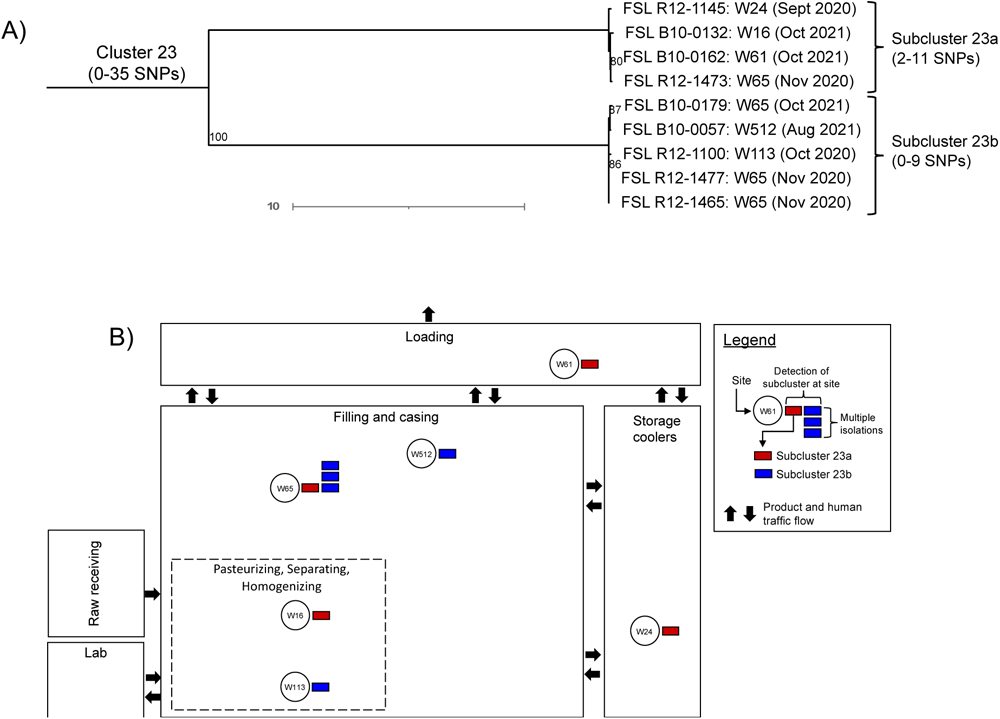
Maximum likelihood phylogenetic tree of plant W *L. seeligeri* cluster 23 and its subclusters (A) and map of the location in which subclusters were identified in the plant (B). For (A), each node represents isolate ID (i.e., FSL R12-1145), site number (i.e., W24), and date (month and year) in which a given isolate was obtained. hqSNP data from cluster 23 was used to create the maximum likelihood phylogenetic tree using RAxML (v 8.2.12) with 1,000 bootstraps and the GTRCAT nucleotide substitution model. Only bootstrap values >70% are presented in the tree, and the tree is rooted at its midpoint. The scale at the bottom of the phylogenetic tree indicates genetic distance. For (B), a colored bar indicates isolation of a given subcluster, and multiple bars of the same color are shown for a given site to indicate the number of samplings when a subcluster was isolated.

On the other hand, plant CL *L. innocua* cluster 10 contained 3 subclusters that were highly related, but were isolated <6 months apart (Figure 6A). These subclusters might represent multiple individual re-introduction events of cluster 10 into the plant, as supported by the finding that subcluster 10a was localized in a different area of plant CL compared to subclusters 10b and 10c, and subclusters 10b and 10c were both isolated from site CL22, which is an external door in the storage cooler and thus represents a site which could facilitate the re-introduction of cluster 10 (Figure 6B). Moreover, singleton isolate FSL B10-0243, which showed 73-78 hqSNP differences from all other isolates in cluster 10 (Supplementary Table 3), was obtained from site CL27, a wall port in the raw receiving area that allows access of raw milk from plant CL’s farm into the plant. As plant CL represents one of the two farmstead plants evaluated in this study (Table 1), one might speculate that plant CL’s farm environment may represent a potential source through which cluster 10 was repeatedly re-introduced into the plant.

**Figure 6.**
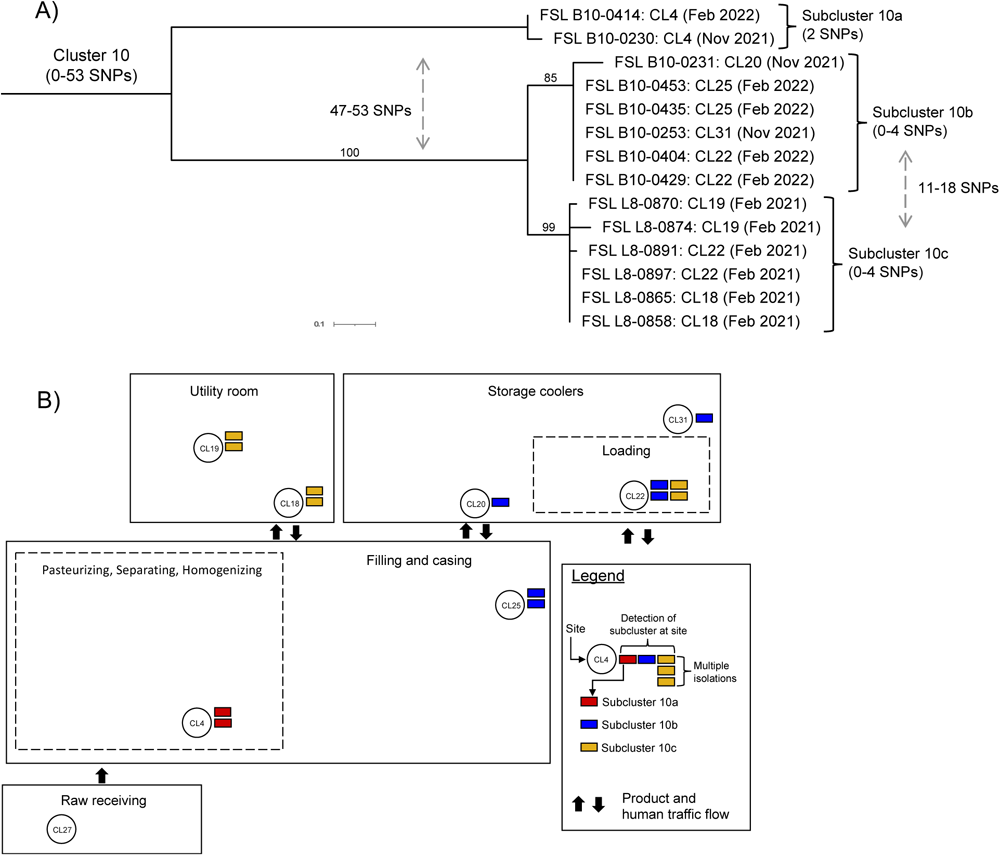
Maximum likelihood phylogenetic tree of plant CL *L. innocua* cluster 10 and its subclusters (A) and map of the location in which subclusters were identified in the plant (B). For (A), each node represents isolate ID (i.e., FSL B10-0414), site number (i.e., CL4), and date (month and year) in which a given isolate was obtained. hqSNP data from cluster 10 was used to create the maximum likelihood phylogenetic tree using RAxML (v 8.2.12) with 1,000 bootstraps and the GTRCAT nucleotide substitution model. Only bootstrap values >70% are presented in the tree, and the tree is rooted at its midpoint. The scale at the bottom of the phylogenetic tree indicates genetic distance. For (B), a colored bar indicates isolation of a given subcluster, and multiple bars of the same color are shown for a given site to indicate the number of samplings when a subcluster was isolated.

### Pre-operation sampling in conjunction with mid-operation sampling can yield valuable insights regarding practices that contribute to *Listeria* persistence

For both initial and follow-up sampling events, there were 42 instances across six plants (i.e., plants CL, CN, CO, CP, CQ, and N) in which the same site tested positive for the same *sigB* AT of *Listeria* at both pre- and mid-operation sampling times. hqSNP analysis was performed to further compare the relatedness of the paired pre- and mid-operation isolates collected in these 42 instances, which revealed that, in nearly all (41/42, 98%) instances, these paired pre- and mid-operation isolates were highly related (i.e., ≤10 hqSNP differences) (Table 7). Additionally, in over half of these instances (23/42, 55%) these paired isolates were nearly identical (i.e., 0-2 hqSNPs) from one another. Notably, all sites where highly related isolates were obtained both pre- and mid-operation represented Zones 3 (*n*=23) and 4 (*n*=18), with structural ontologies of drains (*n*=16), floors (*n*=7), doorway thresholds (*n*=5), wheels (*n*=4), doors (*n*=3), equipment legs (*n*=2), squeegees (*n*=2), and boots and forklifts (both *n*=1). These results suggest that cleaning and sanitation procedures were ineffective at controlling *Listeria* in several SMDPs, particularly at sites representing drain and floor structures, which may represent areas at higher risk for *Listeria* persistence. For example, the 4 drain sites in plant CO that showed highly related *L. newyorkensis* isolates both pre- and mid-operation in one sampling event were represented in cluster 25 (Table 7), and all isolates represented in cluster 25 showed 0-5 hqSNP differences from one another, were obtained drains and floors exclusively, and were isolated over >11 months (Figure 7A).

**Figure 7.**
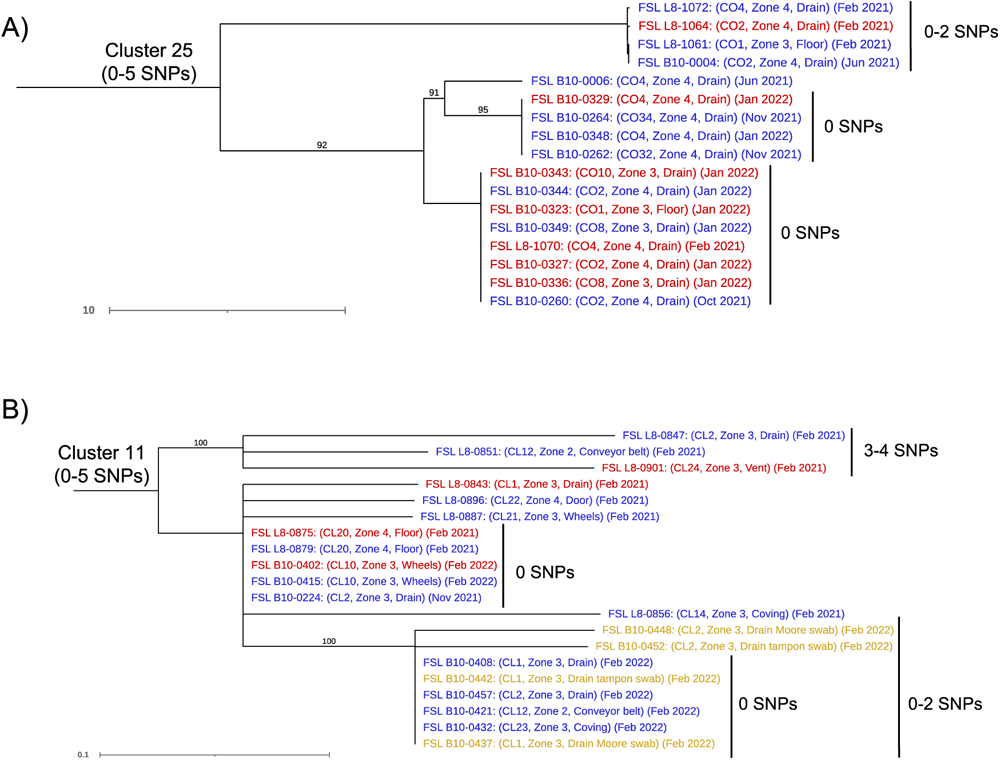
Maximum likelihood phylogenetic trees of hqSNP clusters 25 (plant CO, *L. newyorkensis*) (A) and 11 (plant CL, *L. innocua*) (B) that showed persistence/re-introduction over >6 months. For each node, isolate ID (i.e., FSL L8-1072), site number (i.e., CO4), Zone, site structure, and date (month and year) in which a given isolate was obtained are indicated. Isolates obtained from environmental sponges collected pre-operation (i.e., after cleaning and sanitation but before the next production cycle) are indicated in red; isolates obtained from environmental sponges collected mid-operation (i.e., at least 4 h into a given production cycle) are indicated in blue; isolates obtained from Moore or tampon swabs are indicated in yellow. hqSNP data from clusters were used to create maximum likelihood phylogenetic trees using RAxML (v 8.2.12) with 1,000 bootstraps and the GTRCAT nucleotide substitution model. Only bootstrap values >70% are presented in trees, and trees are rooted at their midpoint. The scale at the bottom of each phylogenetic tree indicates genetic distance.

**Table 7.** hqSNP differences among *Listeria* isolates obtained from the same site at both pre- and mid- operation sampling times in either the initial or follow-up sampling event.

| Sampling event | Site | Zone | Site structure<sup>a</sup> | No. of hqSNP differences | Species (hqSNP cluster isolates are represented in<sup>b</sup>) |
| --- | --- | --- | --- | --- | --- |
| **Plant CL** |  |  |  |  |  |
| Initial | CL18 | 4 | Floor | 0 | *L. innocua* (10) |
| Initial | CL19 | 4 | Floor | 4 | *L. innocua* (10) |
| Initial | CL20 | 4 | Floor | 0 | *L. innocua* (11) |
| Initial | CL22 | 4 | Door | 1 | *L. innocua* (10) |
| Follow-up | CL10 | 3 | Table wheels | 0 | *L. innocua* (11) |
| Follow-up | CL22 | 4 | Door | 0 | *L. innocua* (10) |
| Follow-up | CL25 | 3 | Squeegee | 0 | *L. innocua* (10) |
| **Plant CN** |  |  |  |  |  |
| Initial | CN1 | 3 | Doorway threshold | 5 | *L. innocua* (13) |
| Initial | CN2 | 3 | Drain | 2 | *L. innocua* (13) |
| Initial | CN4 | 3 | Cart wheels | 7 | *L. innocua* (13) |
| Initial | CN9 | 3 | Drain | 5 | *L. innocua* (13) |
| Initial | CN12 | 3 | Equipment leg | 4 | *L. innocua* (13) |
| Initial | CN13 | 3 | Floor | 2 | *L. innocua* (13) |
| Initial | CN15 | 4 | Floor | 0 | *L. innocua* (13) |
| Initial | CN16 | 3 | Pallet jack wheels | 0 | *L. innocua* (13) |
| Initial | CN17 | 4 | Drain | 5 | *L. innocua* (13) |
| Initial | CN18 | 4 | Drain | 9 | *L. innocua* (13) |
| Initial | CN19 | 3 | Doorway threshold | 7 | *L. innocua* (13) |
| Initial | CN20 | 4 | Doorway threshold | 6 | *L. innocua* (13) |
| Initial | CN21 | 4 | Drain | 2 | *L. innocua* (13) |
| Initial | CN22 | 4 | Drain | 7 | *L. innocua* (13) |
| Initial | CN23 | 4 | Doorway threshold | 4 | *L. innocua* (13) |
| Initial | CN25 | 3 | Cart wheels | 3 | *L. innocua* (13) |
| Follow-up | CN22 | 4 | Drain | 3 | *L. innocua* (13) |
| **Plant CO** |  |  |  |  |  |
| Initial | CO4 | 4 | Drain | 3 | *L. newyorkensis* (25) |
| Follow-up | CO2 | 3 | Drain | 0 | *L. newyorkensis* (25) |
| Follow-up | CO4 | 4 | Drain | 0 | *L. newyorkensis* (25) |
| Follow-up | CO8 | 3 | Drain | 0 | *L. newyorkensis* (25) |
| **Plant CP** |  |  |  |  |  |
| Initial | CP6 | 3 | Drain | 0 | *L. monocytogenes* (2) |
| Follow-up | CP1 | 3 | Drain | 2 | *L. innocua* (15) |
| Follow-up | CP5 | 3 | Drain | 0 | *L. innocua* (15) |
| Follow-up | CP7 | 3 | Drain | 0 | *L. innocua* (15) |
| Follow-up | CP8 | 3 | Drain | 0 | *L. innocua* (15) |
| Follow-up | CP15 | 3 | Equipment leg | 3 | *L. innocua* (15) |
| Follow-up | CP35 | 3 | Squeegee | 2 | *L. innocua* (15) |
| Follow-up | CP36 | 4 | Floor | 521 | *L. innocua* (15) |
| Follow-up | CP42 | 4 | Door | 1 | *L. innocua* (15) |
| Follow-up | CP45 | 3 | Forklift | 0 | *L. innocua* (15) |
| **Plant CQ** |  |  |  |  |  |
| Initial | CQ24 | 3 | Doorway threshold | 9 | *L. seeligeri* (20) |
| Follow-up | CQ23 | 3 | Boots | 2 | *L. monocytogenes* (3) |
| **Plant N** |  |  |  |  |  |
| Follow-up | N43 | 4 | Floor | 7 | *L. monocytogenes* (5) |
| Follow-up | N43 | 4 | Floor | 10 | *L. innocua* (17) |

<sup>a</sup> Site structure was identified based on ontology analysis of site descriptions as described in Feng et al. (2023).
<sup>b</sup> Refers to hqSNP cluster no. (see Table 6).

Overall, our findings here support that increased emphasis should be placed on effectively cleaning and sanitizing areas of food processing environments outside of food contact and food-contact adjacent sites. This is particularly important, given that the dynamic nature of a food processing environment can facilitate the transfer of persistent *Listeria* from lower risk sites to high-risk sites, as is also supported by our data. For example, in plant CL, two lower risk (i.e., Zone 3 or 4) sites representing floor (i.e., site CL20) and wheel (i.e., site CL10) structures showed the presence of cluster 11 both pre- and mid-operation within one sampling event (Table 7), and isolates classified in this same cluster were also obtained from environmental sponge samples collected at a Zone 2 conveyor belt on a fluid milk filler (i.e., site CL12) when the site was swabbed mid-operation, but not pre-operation, in both initial (i.e., FSL L8-0851) and follow-up (i.e., FSL B10-0421) sampling events (Figure 7B). These results suggest movement of cluster 11 in plant CL’s processing environment during production. Based on this data, we hypothesize that plant CL’s inability to control *Listeria* in several Zones 3 and 4 sites could have contributed to the re-introduction of cluster 11 at site CL12, which further underscores the importance of ensuring the efficacy of cleaning and sanitation procedures across all areas of a food production environment.

### SSI-1 and PMSCs in *inlA* were identified in over 50% of LM isolates sequenced

All 116 LM isolates represented in clusters and singletons were characterized through *in silico* MLST subtyping, which revealed that these isolates represented 2 lineages (i.e., lineage I and II), 10 unique clonal complexes (CC), and 11 unique sequence types (ST) (Figure 8). Most LM represented lineage II (*n*=85) compared by lineage I (*n*=31), and the top three CC and ST identified included CC9 (all ST1113, *n*=54), CC5 (all ST3189 *n*=27), and CC11 (*n*=19), which represented both ST371 (*n*=17) and ST11 (*n*=2). Two isolates (i.e., FSL B10-0031 and FSL R12-1459) did not contain *bglA*, and thus ST could not be assigned; both isolates represented lineage II and CC199 (Figure 8).

**Figure 8.**
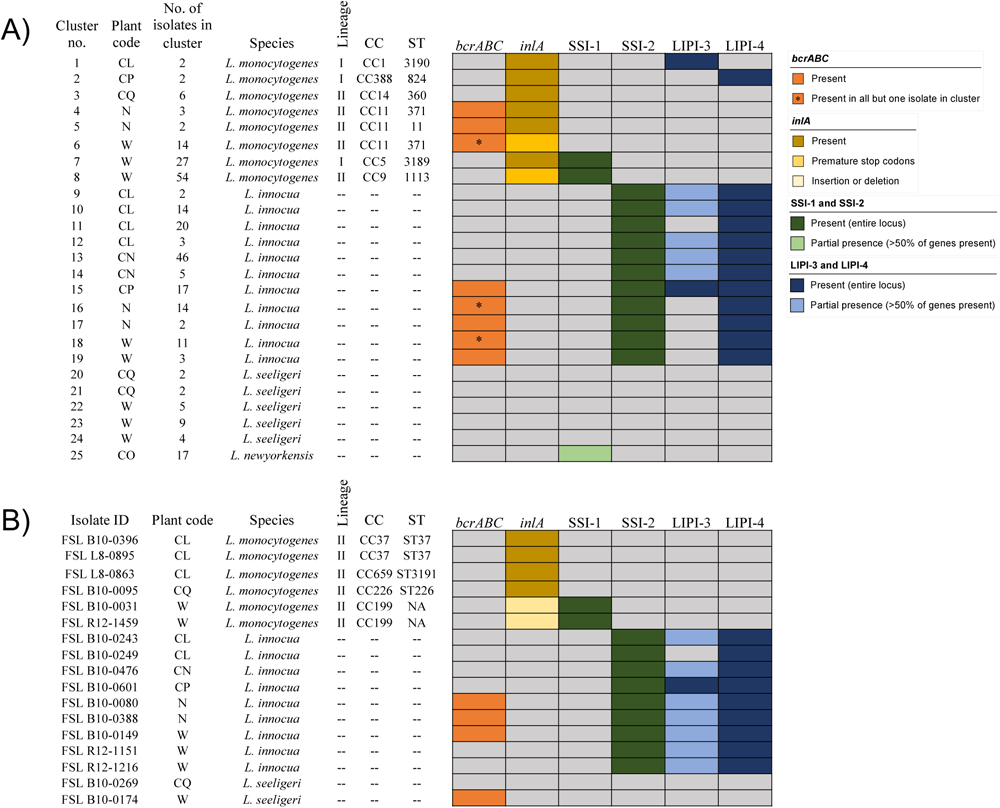
Genomic characteristics of *Listeria* hqSNP clusters (A) and singletons that were not represented in hqSNP clusters (B), including *sigB* allelic type (*sigB* AT), lineage, clonal complex (CC), sequence type (ST), and presence or absence of select genes (i.e., *bcrABC*, *inlA*) and gene loci (i.e., SSI-1, SSI-2, LIPI-3, LIPI-4). ST is reported as “NA” (non-applicable) for singleton isolates FSL B10-0031 and FSL R12-1459 as both isolates did not show the presence of *bglA*, and thus ST could not be assigned.

All LM (*n*=116), *L. innocua* (*n*=146), *L. seeligeri* (*n*=24), and *L. newyorkensis* (*n*=17) genome assemblies represented in clusters and singletons were further screened for the presence of select virulence and stress response associated genes and loci, including *Listeria* pathogenicity islands LIPI-3 and LIPI-4, and stress survival islands SSI-1 and SSI-2 (Figure 8). Interestingly, the presence of both LIPI-4 and SSI-2 appeared to be species specific, with all 146 *L. innocua* isolates showing full matches (>90% similarity over >90% of gene length) to both LIPI-4 and SSI-2, while no LM, *L. seeligeri*, or *L. newyorkensis* isolates sequenced showed either full or partial matches to any LIPI-4 or SSI-2 genes. Conversely, the presence of LIPI-3 and SSI-1 were identified across multiple species, with select isolates showing partial matches to at least 50% of genes associated with these loci. For LIPI-3, a total of 20 isolates, representing *L. innocua* (18/146, 12%) and LM (2/116, 2%), showed full matches to all eight LIPI-3 genes, and 77/146 (53%) of *L. innocua* isolates showed partial presence of the LIPI-3 locus, in which all isolates showed full matches to *llsAGHX*. For SSI-1, 83/116 (70%) LM isolates showed full matches to all five SSI-1 genes, and all 17 *L. newyorkensis* isolates showed full matches to all genes located on the SSI-1 operon except lmo0448. Importantly, nearly all (98/100) *Listeria* isolates that showed full or partial presence of SSI-1 were represented in plant W clusters 7 (*n*=27) and 8 (*n*=54), and plant CO cluster 25 (*n*=17), all of which showed persistence/re-introduction over > 6 months (Table 6).

Additionally, *inlA* was screened for the presence of PMSCs and indels that would confer frameshift mutations in *Listeria* isolates. Overall, as expected, all 116 LM isolates carried *inlA*, and no *L. innocua*, *L. seeligeri*, or *L. newyorkensis* isolates showed either a full or partial match to *inlA*. While *inlA* was present in all LM isolates sequenced, nonsynonymous mutations in *inlA* conferring InlA PMSCs were identified in 68/116 (59%) of LM isolates, representing all isolates from cluster 6 (*n*=14) and cluster 8 (*n*=54), both of which represented isolates collected from plant W over >6 months (Table 6, Figure 8). Additionally, 2 LM singletons (i.e., FSL B10-0031 and FSL R12-1459) from plant W showed the presence of a single frameshift nucleotide deletion in *inlA* (i.e., PMSC mutation type 4) that confers truncation of InlA (Orsi et al., 2007; Van Stelten et al., 2010).

### The presence of *bcrABC* among *Listeria* isolates was correlated with in-plant use of quaternary ammonium compounds

While isolates were also screened for several metal and sanitizer tolerance genes, including *bcrABC*, *emrE*, *qacA*, *qacH*, and *cadAC*, only *bcrABC*, a plasmid-borne sanitizer tolerance cassette, was detected among isolates represented in this study. A total of 67/303 (22%) isolates showed full matches to all three genes on the *bcrABC* cassette, representing *L. innocua* (*n*=48), LM (*n*=18) and *L. seeligeri* (*n*=1) (Figure 8). Interestingly, *Listeria* isolates carrying *bcrABC* represented the majority of isolates sequenced from plants N (22/23, 96%) and CP (17/20, 85%). As these same plants also reported using a quaternary ammonium compound (quat)-based sanitizer for some in-plant sanitation activities (Table 1), these findings suggest that selective pressure through exposure to quats may have played a role in the maintenance of *bcrABC* among *Listeria* in these plants. Meanwhile, less than half of the isolates sequenced from plant W carried *bcrABC* (28/133, 21%). Plant W did not report using a quat-based sanitizer for plant sanitation throughout the 1-year study duration (Table 1), indicating that *Listeria* isolates carrying *bcrABC* can be present in food processing environments in the absence of quat selective pressure.

While all isolates in 5 hqSNP clusters (i.e., clusters 4, 5, 15, 17, and 19) carried *bcrABC*, clusters 6, 16 and 18 each included one isolate where *bcrABC* was not detected (Figure 8). Interestingly, the individual isolate represented in cluster 6 where *bcrABC* was not detected (FSL B10-0052) was nearly identical (0-2 hqSNP differences) to isolates collected ∼6 months before and ∼2 months after FSL B10-0052 was detected in the plant (Figure 9A); a similar trend was also observed for cluster 16 (Figure 9B). For cluster 18, while the individual isolate where *bcrABC* was not detected (FSL B10-0193) was collected in the follow-up sampling event (i.e., Oct 2021), which might suggest loss of *bcrABC* from the overall population, three other isolates obtained in the same sampling did show the presence of *bcrABC* (Figure 9C). Moreover, FSL B10-0193 was closely related (13-18 hqSNP differences) to all other isolates in cluster 18. Overall, these findings indicate that maintenance of *bcrABC* can vary in a given population of closely and even highly related *Listeria* isolates.

**Figure 9.**
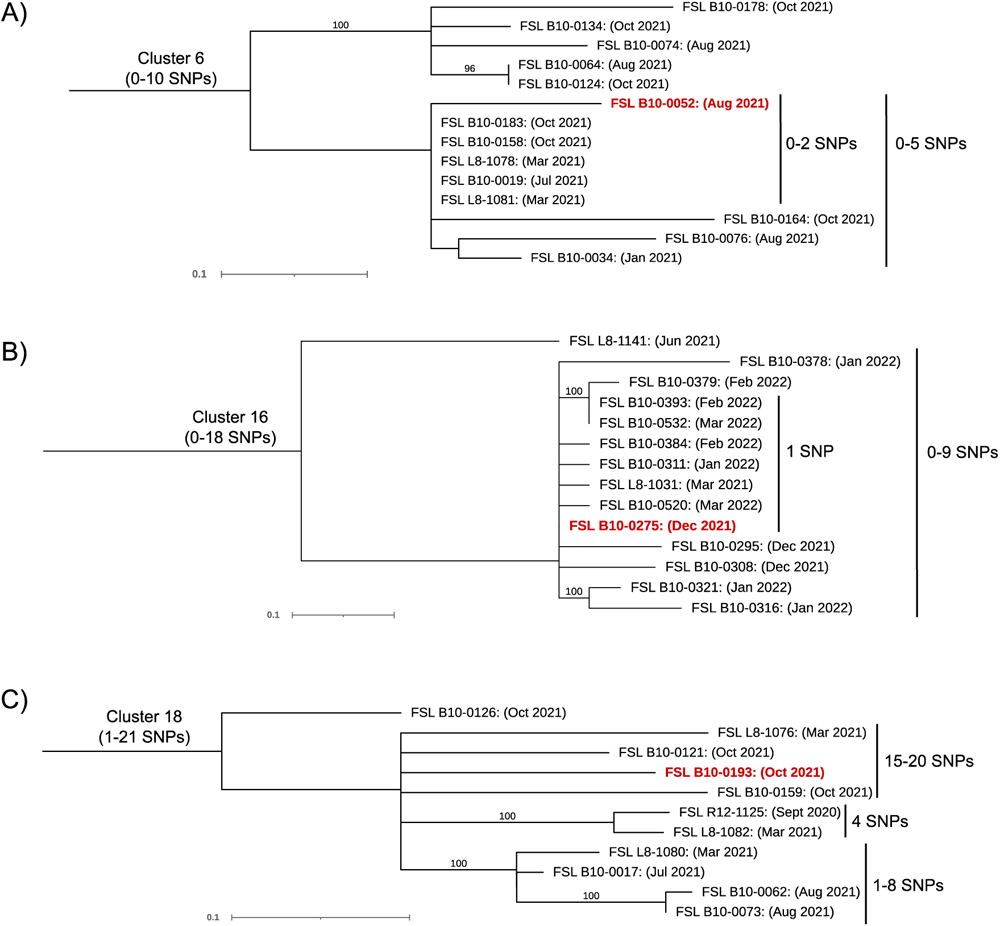
Maximum likelihood phylogenetic trees representing plant W *L. monocytogenes* cluster 6 (A), plant N *L. innocua* cluster 16 (B), and plant W *L. monocytogenes* cluster 18 (C). All clusters contained isolates collected over a period of >6 months, with the majority of isolates carrying the sanitizer tolerance cassette *bcrABC*. For each node, isolate ID (i.e., FSL B10-0052) and the date (month and year) in which a given isolate was obtained are indicated. Isolates within a given cluster that did not carry *bcrABC* are indicated in red. hqSNP data from clusters were used to create maximum likelihood phylogenetic trees using RAxML (v 8.2.12) with 1,000 bootstraps and the GTRCAT nucleotide substitution model. Only bootstrap values >70% are presented in trees, and each tree is rooted at its midpoint. The scale at the bottom of each phylogenetic tree indicates genetic distance.

## DISCUSSION

In this study, we developed and implemented *Listeria* EMPs in eight SMDPs, and evaluated the prevalence and persistence/re-introduction of *Listeria* in these plants over ∼1 year using traditional culture-based detection methods and molecular subtyping (i.e., *sigB* allelic typing, WGS). In addition, we also assessed if simplified environmental sampling strategies, such as collecting environmental sponge samples pre-operation and using passive sampling devices (i.e., Moore and tampon swabs) to collect samples mid-operation, can represent appropriate and effective strategies for detecting *Listeria* in smaller food processing plants. Our findings provide key insights about *Listeria* prevalence, transmission, and persistence in SMDPs, and demonstrate how two simplified environmental sampling strategies, that may be of value to smaller processing plants, can be effectively employed towards *Listeria* EMPs and other control programs.

### Using fixed SNP distance cut-offs to group isolates into SNP clusters may obstruct information that is highly valuable for source tracking in a processing plant

In this study, the 303 putatively persistent *Listeria* isolates characterized through WGS were assigned into initial hqSNP clusters based on a relatively high initial hqSNP distance cut-off (i.e., ≤50 pairwise hqSNP differences between one or more isolates within a given hqSNP cluster). While some studies have reported using a similar strategy (Pettengill et al., 2022; Sullivan et al., 2022; Wang et al., 2018), others with a similar number of isolates have reported using lower SNP distance cut-offs, such as ≤20 or ≤10 SNPs, to assign clusters for further analysis (Andrews et al., 2023; Castro et al., 2021; Lucchini et al., 2023), as they may be more likely to indicate that two or more isolates originated from the same source, e.g., the same plant (Pightling et al., 2018). Importantly, in this study we identified key trends among hqSNP clusters that contained isolates with >21 hqSNP differences that might not have been captured if a lower initial SNP distance cut-off was employed. For example, all isolates in plant CL cluster 10 (hqSNP range: 0-53) were represented in 3 highly related (i.e., ≤10 hqSNP differences) monophyletic subclusters that were isolated <3 months apart, and were either (i) localized in different areas of the plant, or (ii) were identified at sites that could facilitate their introduction into the plant, which may suggest this hqSNP cluster was being re-introduced from a site external to the plant. However, it is important to acknowledge that, alternatively, these 3 subclusters may have evolved from a strain that was introduced in this plant long enough ago to allow for substantial diversification of the cluster.

Even though our relatively high initial hqSNP cut-off was able to capture the relationships between the majority of isolates sequenced (i.e., 286/303, 94%), some isolates that could have provided valuable insights into *Listeria* persistence/re-introduction were still excluded from hqSNP clusters. For example, singleton FSL B10-0243, which showed 73-78 hqSNP differences from all isolates in cluster 10 and thus was excluded from the hqSNP cluster, was isolated from a wall port in the raw receiving area, providing further evidence to suggest the source of cluster 10 in plant CL was associated with plant CL’s farm environment. Overall, our findings indicate that, while fixed SNP distance cut-offs are very useful for rapid decision making, which is especially important for outbreak investigations (Pightling et al., 2018), they might be less appropriate for other cases, such as source tracking in a processing plant or root cause investigations (Alegbeleye & Sant’Ana, 2020), as they might obstruct key information that is highly valuable for identifying root causes of contamination.

### Persistence/re-introduction of *Listeria* is not uncommon in SMDPs, and in some cases may be supported by genetic markers associated with hypovirulence and tolerance to environmental stressors

Of the initial 25 hqSNP clusters identified in this study, over half (14/25, 56%) showed persistence/re-introduction over >6 months, representing five plants. Among these 14 hqSNP clusters, clusters 6 and 8, which represented 40% of CC11 and 100% of CC9 isolates characterized in the study, respectively, showed the presence of *inlA* PMSCs, which have been previously shown to result in a virulence attenuated phenotype in LM isolates (Nightingale et al., 2008). Several previous studies also reported a high frequency *inlA* PMSCs among CC9 isolates associated with food processing environments (Guidi et al., 2021; Maury et al., 2016, 2019). Additionally, we identified the presence of all five genes associated with stress survival islet SSI-1 in 2 LM hqSNP clusters (i.e., clusters 7 and 8) that also showed persistence/re-introduction over > 6 months. Previous studies have also observed a high prevalence of *inlA* PMSCs and SSI-1 among LM showing persistence characteristics in food processing plants (Alvarez-Molina et al., 2021; Kaszoni-Rückerl et al., 2020; Palaiodimou et al., 2021), which may suggest that these genetic markers confer certain phenotypic characteristics that contribute to LM persistence in built environments. For example, presence of *inlA* PMSCs has been reported to be associated with enhanced adherence of LM to polystyrene (Mahoney et al., 2022) and stainless steel surfaces (Piercey et al., 2016) under low temperature storage conditions, and SSI-1 has been reported to enhance LM tolerance to stress conditions such as low pH and high salt concentrations (Ryan et al., 2010). However, these studies were carried out *in vitro*, so there are still knowledge gaps associated with the specific role that *inlA* PMSCs and SSI-1 play in potentially contributing to LM persistence in food processing environments.

In addition, we identified *bcrABC*, a plasmid-borne gene cassette which can confer tolerance of *Listeria* to low-levels of quats (Bolten et al., 2022), in 67/303 (22%) *Listeria* isolates. Previous studies surveying meat and vegetable processing plants (Hurley et al., 2019) and meat and salmon processing plants (Fagerlund et al., 2022) showed similar prevalence of *bcrABC* among *Listeria* populations. Interestingly, 3 hqSNP clusters (i.e., clusters 6, 16, and 18), which showed persistence/re-introduction over >6 months within their respective plants, contained one isolate within each hqSNP cluster that did not carry *bcrABC*. These results suggest that both gain and loss of *bcrABC* can occur among *Listeria* populations showing persistence/re-introduction over time, which is unsurprising given that *bcrABC* is encoded on a plasmid (Dutta et al., 2013), and in general plasmids are highly susceptible to being lost in isolates in the absence of appropriate selective pressures (S. Chen et al., 2017). Importantly, these findings underscore that the presence of *bcrABC*, and similar plasmid-borne genes, might not represent good targets for genetic markers of persistence in food processing environments.

### Virulence pathogenicity islands LIPI-3 and LIPI-4 were more prevalent among *L. innocua* compared to LM

While few WGS-characterized LM isolates showed presence of virulence pathogenicity islands LIPI-3 (1.7%) and LIPI-4 (1.7%), full or partial matches to LIPI-3 and LIPI-4 were identified in many of the *L. innocua* isolates sequenced in this study. In particular, full-length, and partial length (i.e., presence of *llsAGHX*) LIPI-3 was identified in 12% and 53% of *L. innocua* isolates, respectively, and full-length LIPI-4 was identified in all *L. innocua* isolates. While historically more commonly associated with LM, recent efforts to perform more in-depth WGS-based characterizations of non-LM *Listeria* spp. have also identified the frequent presence of LIPI-3 and LIPI-4 in environmental *L. innocua* (Clayton et al., 2014; Lee et al., 2023). Although LIPI-3 and LIPI-4 appear to confer phenotypic characteristics that play an important role in LM hypervirulence (Maury et al., 2016), their functional role in *L. innocua* has not been well studied (Lee et al., 2023). For LIPI-3, one potential hypothesis is that bacteriocin Listeriolysin S (LLS) produced by *llsA*, which has been reported to show the ability to modulate select Gram positive intestinal microbiota during colonization of mammalian hosts (Ribeiro et al., 2023), might exert similar bactericidal effects on microbial communities present in environmental niches (e.g., food processing environments).

### Simplified *Listeria* sampling strategies may be as valuable as traditional sampling strategies for identifying *Listeria* contamination issues, at least in smaller processing plants

In this study, we evaluated how *Listeria* detection patterns compared when collecting environmental sponge samples at the same or similar sites both pre-and mid-operation. Our results showed that, while mid-operation swabbing yielded a slightly numerically higher percentage of *Listeria* positive samples (17%) as compared to pre-operation swabbing (15%), these differences were not significant, suggesting that pre-operation swabbing may represent an alternative sampling approach with comparable efficacy at detecting *Listeria*, at least in SMDPs at early stages of *Listeria* EMP development. Previous studies carried out in plants that manufacture various RTE (Reinhard et al., 2018) and frozen food (Magdovitz et al., 2022) products also reported observing comparable percentages of *Listeria* positive samples across pre-operation and mid-operation samplings. Moreover, one study (Fate et al., 2021) carried out in small to very small RTE meat and poultry plants reported that pre-operational sampling yielded 2-fold higher *Listeria* positive samples compared to mid-operation sampling.

Additionally, for select SMDPs, we also performed a pilot study to evaluate the use of passive sampling devices (i.e., Moore and tampon swabs) for *Listeria* detection in drains mid-operation. Our results showed that Moore and tampon swabs showed similar ability to detect *Listeria* compared to environmental sponges that were also collected mid-operation from the same drain sites, thus suggesting that use of Moore swabs and tampon swabs in drains may represent a possible alternative to mid-operation swabbing with environmental sponges, at least for SMDPs. However, to our knowledge, this is the first report of passive sampling devices being used towards the purpose of *Listeria* detection in a processing plant setting, and thus further studies will be needed to validate this approach.

Overall, our findings indicate that simplified environmental sampling approaches may show similar effectiveness at detecting *Listeria* compared to more traditional approaches, thus indicating that “one-size-fits-all” recommendations for *Listeria* control may be sub-optimal. For example, while current recommendations prescribed by some regulatory agencies for RTE food manufacturing plants to perform environmental sampling mid-operation as part of their *Listeria* EMPs (Canadian Food Inspection Agency, 2023; U.S. Food and Drug Administration, 2017) may be appropriate for larger processing plants with dedicated food safety personnel, they may be less optimal for smaller food processing plants to implement given their limited resources available to devote to EMP related practices (Luber, 2011; Winkler & Freund, 2011). In addition to being potentially more feasible to implement in smaller plants that lack dedicated food safety personnel, both pre-operation sampling and passive sampling techniques can support more straightforward root cause analysis efforts. For example, sites that test positive for *Listeria* pre-operation are more likely to represent contamination sources that are not effectively being controlled by existing cleaning and sanitation procedures (Belias et al., 2022), while sites that test positive mid-operation often represent transient contamination facilitated by the movement of equipment or personnel during the production cycle, and thus may require follow-up sampling (i.e., vector swabbing) to identify the actual primary sites of persistence (Spanu & Jordan, 2020). Moreover, passive sampling devices can capture at least some of the water-mediated movement of *Listeria* within a processing environment over a long period of time (i.e., an entire production cycle) without requiring the same labor input as collecting environmental sponge samples mid-operation. Thus, current recommendations that environmental sampling as part of *Listeria* EMPs should be performed mid-operation exclusively could potentially be re-evaluated, at least for smaller food manufacturing plants.

### Environmental sampling carried out both pre- and mid-operation, in conjunction with WGS data, can support the identification of *Listeria* persistence sites

Here, we identified 41 cases in 5/8 plants in which highly related (i.e., ≤10 hqSNP differences) *Listeria* isolates were identified at the same site both pre- and mid-operation within a single sampling event (i.e., initial or follow-up sampling event). This combination of (i) sampling time metadata and (ii) SNP-based relatedness data indicate that *Listeria* persistence was more likely than re-introduction at these individual sites, with the most probable root cause of this site-specific persistence being ineffective cleaning and sanitation. Importantly, all sites that yielded highly related *Listeria* isolates pre- and mid-operation represented Zones 3 and 4 sites, which suggests that the SMDPs evaluated here may struggle with effectively cleaning and sanitizing sites extending food-contact and food-contact-adjacent surfaces (i.e., Zones 1 and 2); suboptimal sanitary equipment and facility design likely represent an important contributor to this challenge. Given that transfer of *Listeria* from lower risk sites (i.e., Zones 3 and 4) to higher risk sites (i.e., Zone 2) has been documented in previous studies (Sullivan et al., 2022), as well as our study, our results highlight that increased emphasis should be placed on ensuring that Zone 3 and 4 sites are effectively and regularly cleaned and sanitized in SMDPs.

With respect to factors that contribute to *Listeria* persistence in food processing environments, many studies have characterized high-risk sites that could support *Listeria* harborage (Estrada et al., 2020; Pritchard et al., 1995). Here, we identified that the majority of structural ontologies represented among cases of site-specific persistence were at drains and floors, both of which are well-recognized as areas that are often be difficult to clean and sanitize (Simmons & Wiedmann, 2018; Truchado et al., 2022). Notably, we also identified a handful of instances of site-specific persistence at several structures representing moveable equipment, such as cart wheels, squeegees, boots, and forklifts, all of which could potentially facilitate *Listeria* transfer throughout the processing environment (Belias et al., 2022). Overall, these data provide valuable insights about key sites in processing environments that are at high risk for *Listeria* persistence.

### Implementation of *Listeria* EMPs in SMDPs may not yield positive food safety outcomes, unless supported by additional tools and training efforts

Overall, the implementation of *Listeria* EMPs in the SMDPs enrolled in this study resulted in limited reductions in *Listeria* prevalence, as evidenced by the fact that only one plant (i.e., plant CN) showed a significantly lower percentage of *Listeria* positive samples after 1-year of EMP implementation. Therefore, our findings indicate that implementing *Listeria* EMPs using a traditional framework may not yield positive food safety outcomes within the span of 1 year, which differs from previous studies that evaluated the longitudinal implementation of *Listeria* control programs in other small and medium sized food processing plants (Beno et al., 2016; Fate et al., 2021). Based on our findings, we hypothesize that smaller processing plants may need alternative or additional targeted guidance for implementing *Listeria* EMPs. While trainings focused on supporting enhanced education and improving food safety culture in smaller food plants have been previously developed (Evans et al., 2021; Kovacevic, 2022; Magiya, 2023), supplemental trainings focused on the practical implementation of *Listeria* EMPs may prove highly beneficial. Finally, further additional research pursuits focused on identifying and evaluating simplified and size-appropriate sampling strategies, like the ones we preliminary examined in this study, will be valuable to help with informing both future industry and regulatory guidance on this topic.

## ACKNOWLEDGEMENTS

The work presented here was funded by the New York Dairy Promotion Order (Albany, NY) through New York State’s Sponsor Award Number C012388. The authors thank the participating dairy processing plants for their support with environmental sample collection, Dr. Erika Mudrak of the Cornell Statistical Consulting Unit for statistical analyses guidance, and the Institut Pasteur teams for the curation and maintenance of BIGSdb-Pasteur databases (http://bigsdb.pasteur.fr/). The authors also want to thank Karen Ospina of the Cornell Dairy Foods Extension Program for assistance during sampling visits, and Mikayla Henry of the Milk Quality Improvement Program for laboratory support.

## Notes

https://doi.org/10.7298/ah3b-ew05

### Competing Interest Statement

The authors have declared no competing interest.

