## Supplementary Table 3 for "Intensive environmental sampling and whole genome sequence-based characterization of *Listeria* in small and medium sized dairy plants reveal opportunities for simplified and size-appropriate environmental monitoring strategies"

**Table S3.** Characteristics of the 17 singleton *Listeria* isolates that were not represented in hqSNP clusters.

| Isolate ID | Plant code | *sigB* AT^a^ | Month of isolation | Most closely related cluster or singleton | hqSNP range^b^ from most closely related cluster or singleton |
| --- | --- | --- | --- | --- | --- |
| ***L. monocytogenes* singletons** | | | | | |
| FSL B10-0396 | CL | 57 | Feb 2022 | FSL L8-0895 | 80 |
| FSL L8-0895 | CL | 57 | Feb 2021 | FSL B10-0396 | 80 |
| FSL L8-0863 | CL | 57 | Feb 2021 | Cluster 3 | >100^c^ |
| FSL B10-0095 | CQ | 57 | Aug 2021 | Cluster 3 | >100 |
| FSL B10-0031 | W | 57 | Jan 2021 | FSL R12-1459 | 57 |
| FSL R12-1459 | W | 57 | Nov 2020 | FSL B10-0031 | 57 |
| ***L. innocua* singletons** | | | | | |
| FSL B10-0243 | CL | 31 | Nov 2021 | Cluster 10 | 73-78 |
| FSL B10-0249 | CL | 31 | Nov 2021 | Cluster 10 | >100 |
| FSL B10-0476 | CN | 110 | Mar 2022 | Cluster 14 | >100 |
| FSL B10-0601 | CP | 11 | Mar 2022 | Cluster 15 | >100 |
| FSL B10-0080 | N | 37 | Jul 2021 | FSL B10-0388 | >100 |
| FSL B10-0388 | N | 37 | Feb 2022 | FSL B10-0080 | >100 |
| FSL B10-0149 | W | 37 | Oct 2021 | FSL B10-0388 | >100 |
| FSL R12-1151 | W | 37 | Sept 2020 | FSL B10-0080 | >100 |
| FSL R12-1216 | W | 37 | Sept 2020 | FSL B10-0388 | >100 |
| ***L. seeligeri* singletons** | | | | | |
| FSL B10-0269 | CQ | 12 | Nov 2021 | Cluster 20 | >100 |
| FSL B10-0174 | W | 3 | Oct 2021 | Cluster 23 | >100 |

^a^ *sigB* allelic type.

^b^ If only one value is provided for the hqSNP range, only one SNP difference exists.

^c^ Indicates that isolates showed >100 SNP differences from all other isolates in the cluster based on kSNP3 analysis.
